# Sequential transcriptional waves and NF-κB-driven chromatin remodeling direct drug-induced dedifferentiation in cancer

**DOI:** 10.1101/724740

**Authors:** Yapeng Su, Chunmei Liu, Xiang Lu, Guideng Li, Shiqun Shao, Yan Kong, Jihoon W. Lee, Rachel H. Ng, Stephanie Wong, Lidia Robert, Charles Warden, Victoria Liu, Jie Chen, Zhuo Wang, Guangrong Qin, Yin Tang, Hanjun Cheng, Alphonsus H. C. Ng, Songming Peng, Min Xue, Dazy Johnson, Yu Xu, Jinhui Wang, Xiwei Wu, Ilya Shmulevich, Qihui Shi, Raphael Levine, Antoni Ribas, David Baltimore, Jun Guo, James R. Heath, Wei Wei

**Affiliations:** Institute for Systems Biology, Seattle, WA 98109, USA; Division of Chemistry and Chemical Engineering, California Institute of Technology, Pasadena CA 91125, USA; Division of Biology and Biological Engineering, California Institute of Technology, Pasadena CA 91125, USA; Department of Molecular and Medical Pharmacology, University of California – Los Angeles, Los Angeles CA 90095, USA; The Key Laboratory of Carcinogenesis and Translational Research (Ministry of Education), Department of Renal Cancer and Melanoma, Peking University Cancer Hospital and Institute, Beijing, China; Department of Medicine, University of California – Los Angeles, Los Angeles CA 90095, USA; Department of Molecular and Cellular Biology, City of Hope, Duarte, CA 91010, USA; Key Laboratory of Systems Biomedicine (Ministry of Education), Shanghai Center for Systems Biomedicine, Shanghai Jiao Tong University, Shanghai, China; Fudan University Shanghai Cancer Center, Shanghai, China; Key Laboratory of Medical Epigenetics and Metabolism, Institute of Biomedical Sciences, Fudan University, Shanghai, China; The Fritz Haber Research Center, The Hebrew University, Jerusalem, Israel; Department of Surgery, University of California – Los Angeles, Los Angeles, CA 90095, USA; Jonsson Comprehensive Cancer Center, University of California – Los Angeles, Los Angeles, CA 90095, USA

## Abstract

Drug-induced dedifferentiation towards a drug-tolerant persister state is a common mechanism cancer cells exploit to escape therapies, posing a significant obstacle to sustained therapeutic efficacy. The dynamic coordination of epigenomic and transcriptomic programs at the early-stage of drug exposure, which initiates and orchestrates these reversible dedifferentiation events, remains largely unexplored. Here we employ high-temporal-resolution multi-omics profiling, information-theoretic approaches, and dynamic system modeling to probe these processes in *BRAF*-mutant melanoma models and patient specimens. We uncover a hysteretic transition trajectory of melanoma cells in response to oncogene inhibition and subsequent release, driven by the sequential operation of two tightly coupled transcriptional waves, which orchestrate genome-scale chromatin state reconfiguration. Modeling of the transcriptional wave interactions predicts NF-κB/RelA-driven chromatin remodeling as the underlying mechanism of cell-state dedifferentiation, a finding we validate experimentally. Our results identify critical RelA-target genes that are epigenetically modulated to drive this process, establishing a quantitative epigenome gauge to measure cell-state plasticity in melanomas, which supports the potential use of drugs targeting epigenetic machineries to potentiate oncogene inhibition. Extending our investigation to other cancer models, we identify oxidative stress-mediated NF-κB/RelA activation as a common mechanism driving cellular transitions towards drug-tolerant persister states, revealing a novel and pivotal role for the NF-κB signaling axis in linking cellular oxidative stress to cancer progression.

## Introduction

Cell state transitions, characterized by genome-wide transcriptome alterations, are fundamental to numerous biological processes, including stem cell differentiation, immune cell activation, oncogenesis, and cancer cell adaptation to therapies. In a developmental process, these transcriptome alterations are associated with the reconfiguration of the global chromatin landscape and have been demonstrated to occur through “transcriptional waves” driven by specific transcriptional factors (TFs) and collaborative co-factors^1–3^. Similarly, under therapeutic stress, certain tumor cells can also hijack the developmental process to reversibly dedifferentiate towards stem-like, drug-tolerant persister (DTP) states. Such phenotypic plasticity has been reported in a variety of cancer types under different therapy regimens, enabling tumor cells to survive therapeutic treatment prior to the emergence of genetically resistant derivatives^4–8^. Unlike the traditional model of resistance development, which is based on the selection of preexisting resistant clones, this non-genetic process involves dynamic, drug-induced cell-state changes, recently validated by time-lapse imaging and single-cell lineage tracing^9–11^. Despite extensive analysis of resultant dedifferentiated DTPs in previous studies^4,12–14^, how this dynamic dedifferentiation program is mechanistically initiated under therapeutic stress and whether it is orchestrated by temporally ordered transcriptional waves akin to those in developmental biology, remains unclear. Further questions revolve around which key regulators instruct the sequence and pace of the epigenetic and transcriptomic reprogramming during dedifferentiation. Addressing these fundamental questions can provide critical insights into this reversed developmental process that cancer cells exploit to evade therapies and identify targetable vulnerabilities to arrest this process from its inception.

Here we explore these questions using *BRAF*-mutant melanoma models, a paradigmatic example of adaptive cell-state dedifferentiation following driver oncogene inhibition. Continuous treatment of *BRAF*-mutant melanomas with BRAF inhibitors (BRAFi) or other MAPK inhibitors (MAPKi) can lead to the acquisition of a stem-like drug-tolerant state^5,11,12,15–17^. Notably, upon drug removal, the cells can revert to the original drug-sensitive state. We performed a kinetic series of epigenome and transcriptome analyses following BRAFi treatment and subsequent drug removal, followed by a systems-level multi-omic integration of the forward and reverse transition dynamics. Our results revealed that the transition is driven by the sequential operation of two tightly coupled transcriptional waves. Dynamic system modeling and integrated epigenome-transcriptome analysis further elucidated the critical role of NF-κB-driven chromatin remodeling in initiating and driving the dedifferentiation transition, quantitatively revealing the epigenetic nature of cancer cell plasticity. We further explored the molecular mechanisms associated with the initiation of the adaptive cell state transition towards DTPs across multiple cancer types, underscoring the pivotal role of NF-κB transcriptional activation in mediating these unfavorable cell state changes. These insights provide a foundation for the future development of new therapeutic strategies aimed at disrupting the epigenetic mechanisms that enable cancer cells to evade treatment and persist in a drug-tolerant state.

## Results

### Reversible drug-induced dedifferentiation towards therapy resistance in *BRAF*-mutant melanomas

We previously identified a panel of patient-derived *BRAF-*mutant melanoma cell lines that exhibit phenotypic plasticity, allowing them to adapt to the BRAFi treatment by transitioning into a less differentiated, drug-tolerant state^5,17^. Here, we first selected a representative, highly plastic cell line, M397, to investigate the dynamics of the dedifferentiation transition. Upon continuous treatment with the BRAFi, vemurafenib (VEM), these cells dedifferentiated from a drug-sensitive melanocytic state into a transient, slow-cycling state with an expression profile resembling their neural crest developmental ancestor. This initial transition was followed by a progression towards a mesenchymal state, characterized by elevated expression of epithelial-to-mesenchymal (EMT) transition-related genes such as *TGFBI, CDH2, FN1*, along with down-regulation of TFs essential for melanocyte differentiation, particularly *MITF* and *SOX10* (Fig. 1a–c, and Extended Data Fig. 1a–d)^18,19^. The mesenchymal phenotype is notorious for its resistance to BRAFi-targeted therapies as well as other treatment regimens, including immunotherapy^5,15,16,20,21^.

**Fig. 1.**
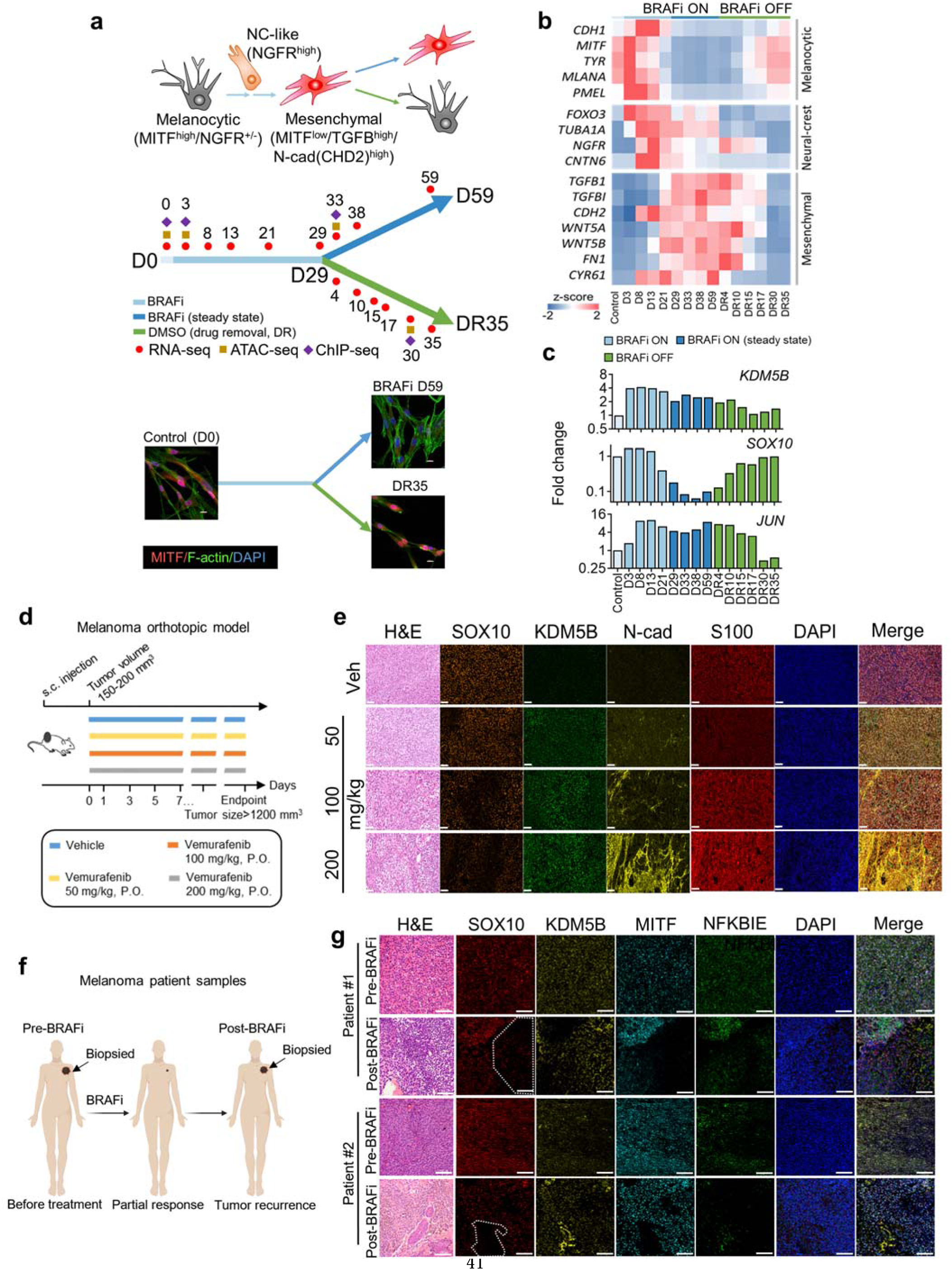
BRAF inhibition-induced dedifferentiation transition towards drug tolerance in melanoma cell lines, mouse xenograft models, and melanoma patient samples. **a.** An illustration of the melanocyte-to-mesenchymal dedifferentiation transition with a transient neural crest state (top) and the experimental timeline (middle). M397 cells were treated with 3 µM BRAFi vemurafenib for 29 days (D29). BRAFi treatment continued for some cells up to D59, while other cells were followed over a 35-day period of drug removal (DR35). Cells were harvested for RNA-seq, ATAC-seq, and ChIP-seq at the specified time points. Immunofluorescent staining (bottom) shows the remodeling of actin filaments and reduction of MITF in mesenchymal-like drug-tolerant cells (D59) and the recovery after drug removal (DR35). **b.** A heatmap showing the expression levels of cell state-specific genes during the dedifferentiation transition and its reversion. **c.** Gene expression levels, normalized to D0, of *KDM5B, SOX10* and *JUN,* over the course of the drug-induced dedifferentiation transition and drug removal, measured by time-series RNA sequencing. **d.** A schematic illustration of the experimental setting for the *BRAF*-mutant melanoma mouse xenograft model. **e.** H&E and mIHC staining with antibodies targeting cell state-associated markers in tumor sections derived from the melanoma mouse xenograft model treated with varying doses of vemurafenib. A dose-dependent loss of SOX10 and increase in KDM5B and N-Cadherin were observed (scale bar: 50 μm). **f.** A schematic illustration of longitudinal tumor biopsy samples collected from patients bearing *BRAF*-mutant melanoma before and after BRAFi treatment. **g.** H&E and mIHC staining of melanoma patient biopsy samples with antibodies targeting cell state-associated markers. The regions highlighted by white dashed lines in the post-BRAFi tissues display reduced expression levels of melanocytic markers MITF and SOX10 and elevated expression of KDM5B, consistent with the molecular characteristics of drug-induced dedifferentiation towards adaptive resistance observed in cell line studies (scale bar: 100 μm).

Consistently, time-series transcriptome analysis revealed a global shift in the transcriptome during the dedifferentiation transition (Fig. 1b, and Extended Data Fig. 1a), with upregulation of several known molecular markers associated with melanoma DTPs, including *KDM5B* and *JUN* (Fig. 1c). There was also an increase in EMT signatures accompanied by the downregulation of MITF targets in the dedifferentiated DTP states (Fig. 1b, Extended Data Fig 1d, and Supplementary Dataset 1)^12,13^. An additional month of drug exposure (Day-29 (D29) to D59) maintained the cells in a state with a relatively stable transcriptome profile, potentially reflecting the drug-resistant mesenchymal state as a stable steady state (i.e. an “attractor”) in the transcriptome space (Fig. 1b and Extended Data Fig. 1a)^6^.

Drug removal (DR) triggered a return to a state that was characterized by a transcriptome profile, cell-cycle characteristics, and BRAFi sensitivity almost identical to the untreated (D0) state (Fig. 1a–c and Extended Data Fig. 1a–f). This reversibility supports a non-genetic mechanism of drug tolerance. We further confirmed these findings in other *BRAF*-mutant melanoma cell lines exhibiting varying degrees of phenotypic plasticity. Upon BRAFi rechallenge after a period of drug removal, the cells exhibited drug sensitivities, cell-cycle characteristics, and drug response profiles similar to those of the untreated cells (Extended Data Fig. 1e,f)^5^. These results demonstrate that phenotypically plastic melanoma cells can reversibly switch between a dedifferentiated DTP state and a differentiated drug-sensitive state upon drug treatment and removal.

### Drug-induced dedifferentiation in melanoma mouse models and patient specimens

To determine whether the drug-induced dedifferentiation transition observed *in vitro* extends to *in vivo* patient-derived tumor models, we established a mouse xenograft model using patient-derived *BRAF*-mutant melanoma cells and treated the mice with BRAFi VEM at a dose escalation setting (Fig. 1d and Extended Data Fig. 2a). Multiplex immunohistochemistry (mIHC) staining displayed dose-dependent transition kinetics, with the most pronounced dedifferentiation signatures—characterized by reduced SOX10 and elevated KDM5B and N-Cadherin—appearing in mice treated with the highest dose (Fig. 1e and Extended Data Fig. 2b). Additionally, we collected longitudinal tumor biopsies from patients with *BRAF*-mutant melanoma who exhibited a partial response to BRAFi. We sampled these biopsies at diagnosis before BRAFi treatment and at the onset of therapy resistance (Fig. 1f). The analysis revealed a spatially heterogeneous response to BRAFi treatment, with lineage-specific and epigenetic-related markers coordinately altered from region to region. Regions showing dedifferentiation signatures characterized by significantly reduced MITF and SOX10 also displayed elevated levels of drug-tolerant marker KDM5B, consistent with cell line studies (Fig. 1g and Extended Data Fig. 2c). Other regions retained similar MITF and SOX10 expression and exhibited reduced levels of KDM5B. Likewise, an analysis of published transcriptome data from longitudinal *BRAF-*mutant melanoma patient samples before and after BRAFi (or MAPKi) treatment revealed analogous dedifferentiation signatures, transitioning from the melanocytic state towards the drug-tolerant mesenchymal state (Extended Data Fig. 2d and Supplementary Dataset 2)^16,22^. Collectively, these results demonstrate the *in vivo* and clinical relevance of a drug-induced cell state dedifferentiation transition in melanoma.

### Information-theoretic analysis reveals sequential transcriptional waves orchestrating a “hysteretic” transition trajectory

To simplify the global transcriptome dynamics, where thousands of genes are altered reversibly upon drug treatment and removal, we hypothesized that the drug-induced dedifferentiation process mirrors its developmental counterpart, with transcriptome dynamics proceeding through a few temporally ordered “transcriptional waves.” To extract these transcriptional waves from the temporal transcriptome data, we applied information theoretic surprisal analysis to the time-course transcriptome data of M397.

Surprisal analysis, originally formulated to understand the dynamics of nonequilibrium systems, has been extended to characterize and simplify complex biological processes^23–28^. It approximates the partition function of molecular species within a cell’s molecular ensemble to assess the maximum entropy of those biomolecules. This method de-convolutes the transcriptome dynamics into a global stable state, plus a series of time-dependent gene modules (Eq. 1, see Methods)^24,29^.

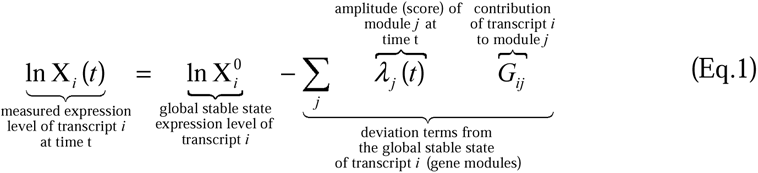

In Eq. 1, X_*i*_^0^ represents the global stable state expression level of transcript *i*. In the summation term on the right side of Eq. 1, the temporal module score, λ*_j_(t),* denotes the weight of gene module *j* at time *t,* while *G_ij_* is the contribution of transcript *i* to module *j*. Using Eq. 1, we de-convoluted the transcriptional dynamics into a global stable state plus two time-varying gene modules.

We visualized the transcriptional stable state and the two modules as self-organizing mosaic (SOM) maps (Fig. 2a) (see Methods)^30^. In these SOMs, the whole transcriptome was assigned to 625 rectangular “tiles” (SOM nodes), each representing a mini-cluster of genes. These tiles form a pattern within a 2-dimensional mosaic map on the SOM grid. Tiles in the same location represent the same group of genes across different conditions, and tiles containing module-specific leading edge genes (genes with large *G_ij_* scores) with similar expression kinetics cluster together as neighboring tiles. When the global stable state and the first two gene modules are summed to predict the whole transcriptome, the resultant SOMs closely recapitulate the experimental data at all the time points, indicating that two time-dependent gene modules are sufficient to capture the transcriptome dynamics of the BRAFi-induced dedifferentiation transition and its reversion (Fig. 2a).

**Fig. 2.**
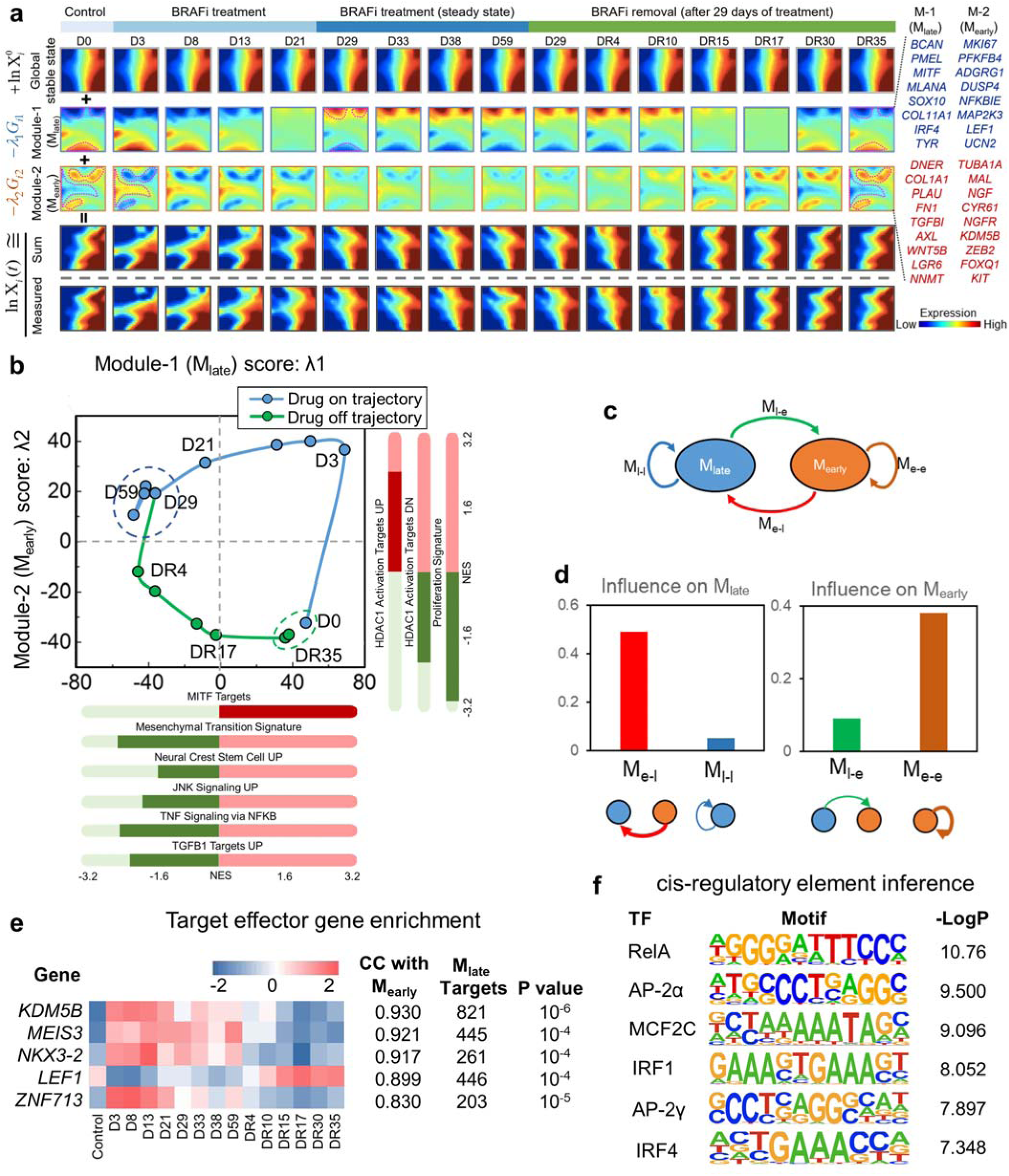
Hysteretic transition trajectory and sequential transcriptional waves during the drug-induced dedifferentiation transition. **a.** Application of surprisal analysis to the time-series transcriptome data over the dedifferentiation transition. The transcriptome data are decomposed into a time-invariant global stable state and two time-dependent gene modules, illustrated as self-organizing maps (SOMs). Adding the expression of the global stable state and first two gene modules recapitulates the experimentally measured transcriptome profiles. Tile clusters containing leading-edge genes associated with each gene module are highlighted by dash lines. Representative module-specific leading-edge genes, whose expression levels coordinately increased (red) or decreased (blue) over the dedifferentiation transition, are listed to the right. **b.** The cyclic trajectory of the dedifferentiation transition plotted in the landscape defined by the first two gene modules. The blue and green dashed lines encircle the milieu of the mesenchymal-like drug-tolerant state and drug-naïve state, respectively. Selected enriched molecular processes (nominal p < 0.05) associated with each gene module are listed. NES: normalized enrichment score. **c.** A schematic illustration of the ODE model for the gene-module interactions. **d.** The module-module interaction coefficients in the ODE model were determined by fitting the model to the average expression levels of genes associated with each gene module. **e.** A list of enriched TFs/co-factors ranked according to their absolute Pearson correlation coefficients (CC) with M_early_ scores, with relative temporal expression levels (z-scores) shown as a heat map. The target gene number and p-value for each enriched element are listed to the right. **f.** Motifs enriched from cis-regulatory elements of M_late_-associated genes. The top six most significantly enriched motifs are listed. -Log (p values) are shown to the right.

We mapped the cell-state transition trajectory by projecting the time-resolved transcriptome data onto a cell state space defined by the first two gene modules, with each axis representing the temporal module scores (Fig. 2b and Supplementary Dataset 3). While previous reports claimed a reversible process for adaptive dedifferentiation transition in melanoma^5,10,15^, our more granular data set revealed a cyclic loop comprised of forward (blue) and reverse (green) trajectories that did not overlap, with two gene modules operating sequentially. Module-2, termed M_early_, was induced first by BRAFi and fully inverted (i.e. the module score changed from negative to positive) by 3 days (D3) of drug treatment, then only slightly decreased in module score over the first month of drug treatment. In contrast, module-1, termed M_late_, displayed sharp changes from D3 to D29, inverting at D21 (Fig. 2b). Neither module changes significantly from D29-D59, suggesting that cells stabilized in a new steady state. Upon drug removal, M_early_ immediately inverted, and then slowly decreased in module score out to DR30. M_late_ displayed consistent increase upon drug removal and inverted at around DR17 (Fig. 2b). These inversion points for M_early_ and M_late_ suggest that two sequential cell-state transition steps separate the drug-sensitive steady state at D0 (or DR30-DR35) and the drug-resistant steady state at D29-D59. Consistently, gene clusters associated with these modules change in sync, reflecting the temporal transcriptional phases along the transition. (Fig. 2a, see clustered tiles circled by dashed lines and representative leading-edge genes listed to the right). The sequential operations of the two modules upon drug exposure and removal suggest the action of two transcriptional waves controlling the cell-state transitions.

The enrichment of module-specific genes revealed molecular signatures and pathways associated with each gene module (Fig. 2b and Supplementary Dataset 4). M_early_ positively associated with HDAC1 activity and negatively associated with cell-cycle regulation, suggesting rapid histone modification and cell cycle arrest upon drug exposure. M_late_ was enriched with gene signatures of neural-crest and mesenchymal lineage specifications, as well as of relevant regulatory pathways, including NF-κB and TGFβ. This is consistent with coordinated changes in lineage-related leading edge genes, such as the reduction of *MLANA* and *PMEL,* and the increase of *TGFBI,* and *FN1* in M_late_ during transitions towards less differentiated states (Fig. 2a,b). These results suggest that the dedifferentiation program associated with M_late_ is triggered after initial chromatin changes associated with M_early_.

A striking result of this analysis is that, for the reversible dedifferentiation process, the forward (drug on) and reverse (drug off) trajectories are not identical but exhibit hysteresis (Fig. 2b). For example, cells at the M_late_ inversion points (D21 for the forward trajectory and DR17 for reverse) exhibit nearly identical M_late_ module scores but are distinguished by a completely inverted M_early_ module (see the SOMs of Fig. 2a at D21 or DR17). This suggests that the cells at these two intermediate transition states possess similar lineage-associated transcriptome profiles but very different chromatin configurations, which are imprinted by their history.

### Dynamic interactions between transcriptional waves reveals early-acting transcription factors initiating the dedifferentiation transition

To discern whether M_early_ and M_late_ operate independently or are coupled, we modeled their dynamics using an ordinary differential equation (ODE) model that resembles a simple two-gene feedback circuit encompassing all possible interactions within a two-body system (Eq. 2 and Fig. 2c, See Methods)^31^.

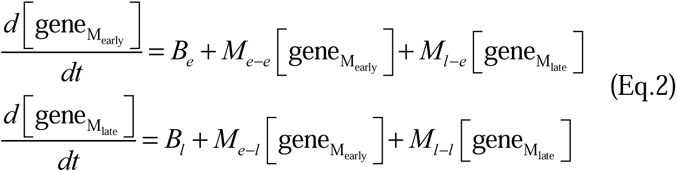

Here, the bracketed terms denote the averaged expression level of module-associated genes at a given time point. *B_e_* and *B_l_*represent the basal production of M_early_- and M_late_-associated genes, respectively. The coefficients *M_e-e_* and *M_l-l_*reflect the self-regulation of M_early_ and M_late_ gene expression, while the *M_e-l_* and *M_l-e_*indicate the regulation of M_late_ gene expression by M_early_ and vice versa.

The best-fitted parameters reveal that M_early_ exerts significant control over both itself and M_late_. In contrast, M_late_ only minimally influences either itself or M_early_. This is evidenced by the much larger coefficients *M_e-e_* and *M_e-l_* compared to *M_l-l_* and *M_l-e_* (Fig. 2d and Extended Data Fig. 3 and Table 1). These results suggest that the transcriptional waves are coupled in a unique way, with TFs/co-factors within M_early_ regulating downstream effector genes not only within M_early_ itself but also within M_late_, thereby driving both the forward and reverse cell state transitions.

To resolve the specific drug-induced TFs/co-factors within M_early_ that drive the M_late_ process, we employed two complementary approaches. First, we assessed TFs/co-factors associated with M_early_ and their downstream targets in M_late_ genes (Extended Data Fig. 4a). Here the association refers to a strong temporal alignment between gene expression levels and module scores (Pearson correlation coefficient |r| > 0.8). We identified several M_early_-associated TFs and co-factors with downstream targets overrepresented in M_late_-associated genes. Some (e.g. LEF1) are known to regulate the expression of melanocyte differentiation effector genes, while others are novel (e.g. MEIS3, NKX3-2) but have been reported to be involved in neural-crest development in other contexts (Fig. 2e)^16,32,33^. Notably, we identified the histone demethylase KDM5B at the top of the list. KDM5B has been reported as a molecular marker associated with DTP state in many cancers including melanoma^13^. However, the mechanistic understanding of how KDM5B-mediated chromatin remodeling leads to drug tolerance in melanoma remains unclear.

The sharp increase of *KDM5B* expression by D3 echoes the enrichment of HDAC1 activity within M_early_ (Figs. 1c and 2b). Consistently, enrichment analysis showed a significant downregulation of KDM5B and HDAC1 target genes at D3 compared to DMSO control, suggesting increased activities of these two histone modifiers to reduce their target activation histone marks, such as H3K4me3 and H3K27ac, and consequently repress their target gene expression (Extended Data Fig. 4c). Together, these findings suggest rapid histone modifications associated with M_early_ activation immediately after drug treatment.

For the second approach, we enriched the cis-regulatory elements in the promoter regions of M_late_- associated genes (Pearson |r| > 0.8) to infer the binding motifs of TFs that can potentially regulate these M_late_ effector genes. (Fig. 2f and Extended Data Fig. 4b). Enriched TFs included AP-2 paralogs, MEF2C, and IRF1/4, consistent with their known role to co-regulate melanocyte differentiation in parallel with MITF and SOX10^34–36^. The most significantly enriched motif was RelA, an NF-κB family member whose activation has been reported as a protective early response to therapeutic stress^37,38^ (Fig. 2f). The high enrichment of the RelA binding motif in lineage-associated M_late_ genes suggests that RelA transcriptional activation is a key early-acting component driving the sequential transcriptional dynamics during dedifferentiation.

### Integrated multi-omics analysis of NF-**κ**B/RelA-driven chromatin remodeling reveals downstream transcription factors driving the late-acting transcriptional wave

We sought to further investigate the roles of early-acting RelA and histone modifiers KDM5B and HDAC1 in driving the transcriptional dynamics of cell-state dedifferentiation in M397. This was achieved by assessing drug-induced changes of genome-wide chromatin accessibility and two activation histone marks, H3K4me3 and H3K27ac, direct targets of KDM5B and HDAC1, respectively (Extended Data Fig. 5a,b). Like transcriptome, the global chromatin accessibility exhibited reversible changes during BRAFi treatment and removal (Extended Data Fig. 6a,b). K-mean clustering of differential cis-regulatory elements revealed a clear chromatin remodeling, with reduced chromatin accessibility in a subset of cis-regulatory elements (Clusters III/IV) upon drug treatment and a reversion to a profile similar to that of untreated cells upon prolonged drug removal (Fig. 3a). Striking reversibility was also observed in the two histone marks with only minor differences in differential peaks between the long-term drug removal state (DR30) and the treatment naïve state (D0) (Fig. 3b). This epigenomic reversibility may underlie the observed transcriptome and phenotypic reversibility during the cell state transition.

**Fig. 3.**
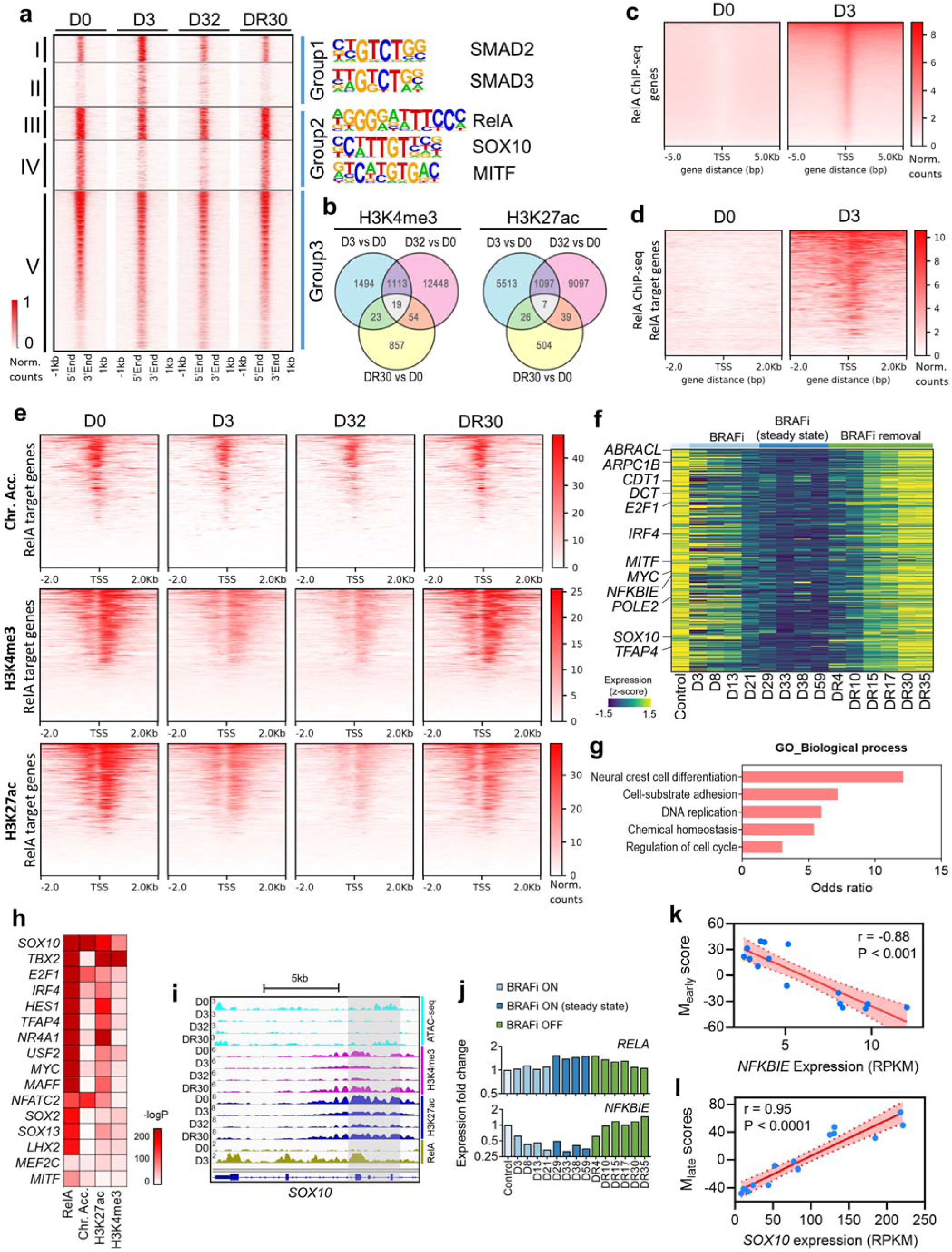
Epigenetic reversibility and NF-κB/RelA-driven chromatin remodeling during the dedifferentiation transition. **a.** Heatmap of chromatin accessibility changes assessed by average ATAC-seq peak signal across all peaks, at selected time points over the transition. K-means clustering of rows identifies five chromatin regions grouped into D3 enriched (group 1), D0/DR30 enriched (group 2), and drug-treatment independent (group 3). Color corresponds to the normalized cis-regulatory element signal. Representative TF binding motifs enriched in the cis-regulatory element clusters with varying accessibility over the transition are listed to the right. **b.** Venn diagrams showing the numbers and overlaps of differential ChIP-seq peaks for H3K4me3 and H3K27ac. Each circle represents changes in those peaks between two time points, while intersections representing shared changes. **c.** Heatmap showing the normalized read counts of genome-wide RelA ChIP-seq peaks in a ± 5kb genomic interval flanking the TSS at different drug treatment time points. **d.** Heatmap showing the normalized read counts of RelA ChIP-seq peaks in a ± 2kb genomic interval flanking the TSS across the 307 RelA target genes at different drug treatment time points. **e.** Heatmaps showing the normalized read counts of chromatin accessibility, H3K4me3, and H3K27ac in a ± 2kb genomic interval flanking the TSS across the 307 RelA target genes at different drug treatment time points. **f.** Heatmap showing the expression levels of the 307 RelA target genes over the course of drug-induced dedifferentiation transition and its reversion. **g.** Representative enriched biological processes of the 307 RelA target genes (p < 0.05). **h.** Heatmap showing the statistical significance (p-values) of epigenetic alterations at the RelA binding sites located in the promoter regions of TFs/co-factors within the 307 RelA target genes. **i.** Genome browser view of ATAC-seq and ChIP-seq profiles at the promoter region of *SOX10*, at selected time points across the dedifferentiation transition and its reversion. **j.** Gene expression levels of *RELA* and *NFKBIE,* normalized to D0, over the course of the drug-induced dedifferentiation transition and drug removal, measured by time-series RNA sequencing. **k, l**. Correlation between *NFKBIE* expression levels and M_early_ scores (k) and SOX10 expression levels and M_late_ scores (l) over the course of the dedifferentiation transition. The Pearson correlation coefficient and p-value are listed. The shaded region denotes the 95% CI of the linear fitting.

To gain more mechanistic insight, we searched for over-represented TF binding motifs in the cis-element clusters displaying varying accessibility over the transition and identified TFs associated with melanocytic/neural crest lineage and TGFβ signaling, such as SOX10, MITF, and SMAD2/3. The highly enriched motifs of cis-element clusters with reduced accessibility upon drug treatment also included the M_early_-associated TF, RelA, reinforcing results inferred from the transcriptome data (Fig. 3a and 2f). Notably, RelA ChIP-seq profiling revealed a sharp increase in genome-wide RelA binding upon 3 days of BRAFi VEM treatment (Fig. 3c). These findings suggest that RelA may play a crucial role in rendering chromatin less accessible at its binding sites through interactions with histone-modifying cofactors KDM5B and HDAC1, thereby reducing H3K4me3 and H3K27ac marks at the early-stage of drug exposure.

To further infer RelA target genes potentially modulated by RelA-mediated chromatin remodeling and transcriptional repression, we compiled an analytical pipeline that integrates time-series multi-omics data of the dedifferentiation transition (Extended Data Fig. 6c, See Methods). Specifically, we focused on genes whose promoter regions exhibited increased RelA binding and overlapping reductions in H3K4me3 and H3K27ac peaks at D3. These genes were further filtered based on the consequent reduction in chromatin accessibility at the promoter regions and significantly decreased expression levels at D32, reflecting that the early (D3) RelA-mediated reduction of these activation histone marks led to sustained reduction of chromatin accessibility and transcriptional repression upon continuous *BRAF* inhibition.

We identified 307 such RelA target genes, whose expression levels showed more than a two-fold decrease at D32 and recovery upon drug removal (Fig. 3d-f and Supplementary Dataset 5). Consistently, the promoter regions of these genes displayed a sharp increase in RelA peaks at D3 and a marked reduction of both H3K4me3 and H3K27ac peaks, along with reduced chromatin accessibility after drug exposure, with recovery upon drug removal (Fig. 3d, e). Enrichment analysis of these genes revealed signatures associated with neural-crest cell differentiation, melanosome, cell cycle regulation, focal adhesion, and TGFβ regulation, as well as targets of melanocytic lineage TFs, such as SOX10 and MITF (Fig. 3g and Extended Data Fig. 6d). This confirms that these RelA target genes, associated with RelA-mediated chromatin remodeling and transcriptional repression, play a crucial role in regulating cell state dedifferentiation towards slow-cycling neural-crest and mesenchymal-like states.

To further pinpoint the downstream TFs epigenetically regulated by RelA that drive the M_late_-associated transcriptional wave, we assessed changes in chromatin accessibility and two activation histone marks at the RelA binding sites within promoter regions of all TFs among the 307 RelA target genes (see Methods). *SOX10* exhibited the most significant changes in both the RelA binding and all three epigenetic profiles, suggesting its potential role as a key downstream effector of RelA in cell state dedifferentiation (Fig. 3h). Indeed, the promoter regions of *SOX10* displayed reduced chromatin accessibility and activation histone marks upon drug treatment, which reverted upon drug removal (Fig. 3i). While *RELA* expression levels remained relatively steady throughout the dedifferentiation transition, the expression levels of its repressor, *NFKBIE* (i.e. iκBε), rapidly decreased upon drug exposure and exhibited a strong association (anti-correlation, r = -0.88) with M_early_ kinetics (Fig. 3j,k). NFKBIE acts as a RelA repressor by sequestering RelA in the cytoplasm, thereby blocking its nuclear translocation and transactivation^39,40^. The reduction of *NFKBIE* at the early stage of drug treatment could lead to RelA nuclear translocation and transcriptional activation. These data, in conjunction with the strong correlation (r = 0.95) between *SOX10* expression levels and M_late_ kinetics (Fig. 3l), underscore the pivotal role of RelA (early-acting TF) and SOX10 (downstream TF) in driving the two sequential transcriptional waves associated with the melanoma cell-state regression towards resistance.

### Mechanistic hypothesis and experimental validation of the ROS-mediated NF-**κ**B-driven chromatin remodeling initiating the dedifferentiation transition

The regulatory insights inferred from sequential transcriptional dynamics and temporal epigenome profiling suggested a mechanistic regulatory network (Fig. 4a). We hypothesized that BRAF inhibition disrupts redox homeostasis, leading to a rapid reduction of NFKBIE levels and consequent release of RelA to the nucleus. RelA then recruits KDM5B and HDAC1 to repress the expression of *SOX10*, along with other target genes, by erasing activation histone marks and reducing chromatin accessibility in their promoter regions (Fig. 3e, i). The reduction of *SOX10* expression has been reported to promote melanoma dedifferentiation towards a mesenchymal DTP state through downregulating *MITF* expression and up-regulating TGFβ signaling^12,15,18^. Our hypothesis, if validated, will uncover the mechanistic pathway initiating the dedifferentiation process, leading to *SOX10* repression following BRAFi exposure.

**Fig. 4.**
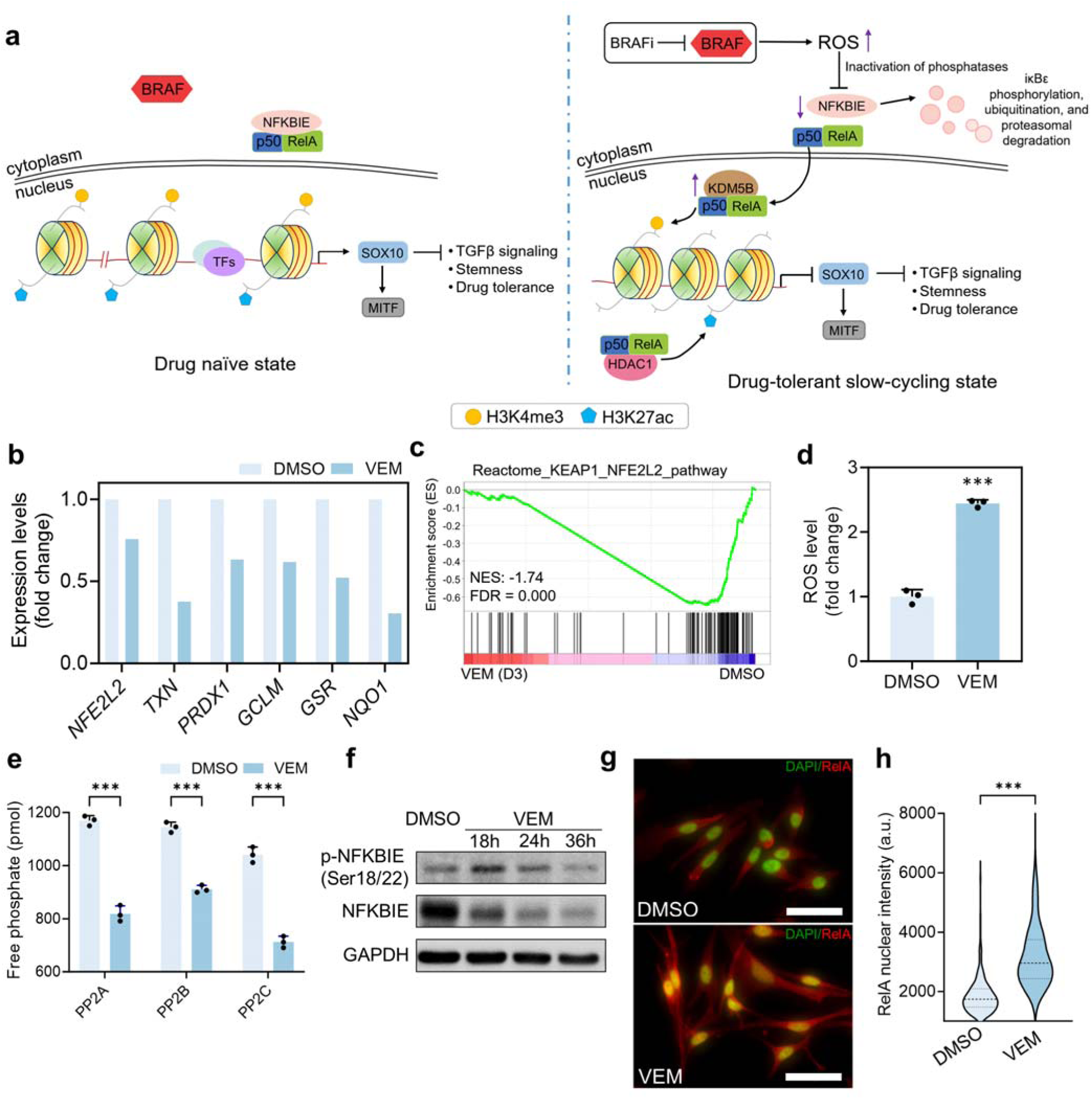
Oxidative stress-mediated RelA nuclear translocation at the early stage of drug-induced dedifferentiation transition in M397 cells. **a.** Illustration of the dedifferentiation-associated mechanistic pathway before and after BRAF inhibition. Left panel: retention of RelA in the cytoplasm and open chromatin at the promoter region of *SOX10*. Right panel: BRAF inhibition leads to elevated ROS, promoting NFKBIE phosphorylation and degradation. The reduction of NFKBIE further promotes the translocation of RelA into the nucleus, allowing RelA to recruit histone modifiers KDM5B and HDAC1 to the target genes, reducing chromatin accessibility and epigenetically repressing of *SOX10*. Functional consequences, such as reduced MITF, increased TGFβ signaling, and transition towards less-dedifferentiated drug-tolerant states, result in turn. **b.** Changes in expression levels of critical genes associated with antioxidant defense after 3 days of 3 μM BRAFi vemurafenib (VEM) treatment compared to the DMSO control. **c.** Gene set enrichment analysis showing the significant downregulation (negative enrichment) of NRF2-related pathway genes upon 3 days of VEM treatment. NES: normalized enrichment score. **d.** Change in cellular ROS levels upon 3 days of 3 μM VEM treatment relative to the DMSO control (mean ± SD, ***p<0.0005, n=3). **e.** Changes in protein phosphatase PP2A, PP2B, and PP2C levels upon VEM treatment relative to the DMSO control, with treatment conditions specified in (**d**) (mean ± SD, ***p<0.0005, n=3). **f.** Immunoblotting of phospho(p)-NFKBIE (Ser18/22) and NFKBIE upon DMSO and VEM (3 μM) treatments for specified durations. **g.** Representative fields of view of immunofluorescence staining showing the increased RelA nuclear translocation upon VEM treatment relative to the DMSO control, with treatment conditions specified in (**d**) (scale bar: 50 μm). **h.** Quantitation of RelA nuclear fluorescent intensities of M397 cells upon VEM treatment relative to the DMSO control, with treatment conditions specified in (**d**) (Two-tailed Mann-Whitney test, ***p<0.0005, n=360 fields of view).

To validate this mechanistic hypothesis, we first assessed expression of genes pivotal to cellular antioxidant defense. Notably, all these genes displayed a reduced expression in M397 upon 3 days of VEM treatment (Fig. 4b). This observation, further supported by the significant downregulation of NRF2 pathway enrichment at D3 — a central pathway for protecting cells from oxidative stress—indicates a dysregulated antioxidant response upon oncogene inhibition (Fig. 4c). Consistently, reactive oxygen species (ROS) levels in M397 cells significantly increased following 3 days of VEM treatment (Fig. 4d). ROS accumulation can activate the NF-κB signaling axis by directly phosphorylating iκB or inactivating serine/threonine phosphatases, leading to iκB phosphorylation, ubiquitination, and proteasomal degradation, thereby reducing NFKBIE (iκBε) expression and consequently promoting RelA nuclear translocation^41,42^. Consistently, we found reduced serine/threonine phosphatases activities, elevated NFKBIE phosphorylation and degradation, and increased RelA nuclear translocation and genome-wide binding within the first 3 days of drug exposure (Fig. 3c and Fig. 4e–h). Notably, the reduction of NFKBIE was also observed in *BRAF*-mutant patient tumor specimens post-BRAFi therapy (Fig. 1g and Extended Data Fig. 2c,d), validating the role of cellular oxidative stress in mediating RelA transcriptional activation.

We then quantitatively assessed the binding profiles of RelA, epigenetic co-factors, and relevant histone marks at the *SOX10* promoter regions in M397 cells using chromatin immunoprecipitation followed by quantitative PCR (ChIP-qPCR). BRAF inhibition resulted in elevated binding of RelA, KDM5B, and HDAC1 to the *SOX10* promoter, along with a consequent decrease in H3K4me3 and H3K27ac histone marks and reduced chromatin accessibility at the same site as soon as within 3 days (Figs. 3i and 5a). These drug-induced changes persisted as cells transitioned to DTPs by D32 (Fig. 5a), leading to reduced *SOX10* expression levels (Fig. 1c). Upon drug removal, these binding profiles reverted to the levels observed in untreated cells (Fig. 5a). Co-immunoprecipitation confirmed the interaction of RelA with KDM5B and HDAC1, suggesting that RelA can form a complex with these histone-modifying enzymes (Fig. 5b). These findings support the role of RelA in recruiting histone-modifying enzymes as co-factors to remodel the chromatin and epigenetically repress the expression of *SOX10*, thereby mediating the expression changes of downstream lineage-associated effectors.

**Fig 5.**
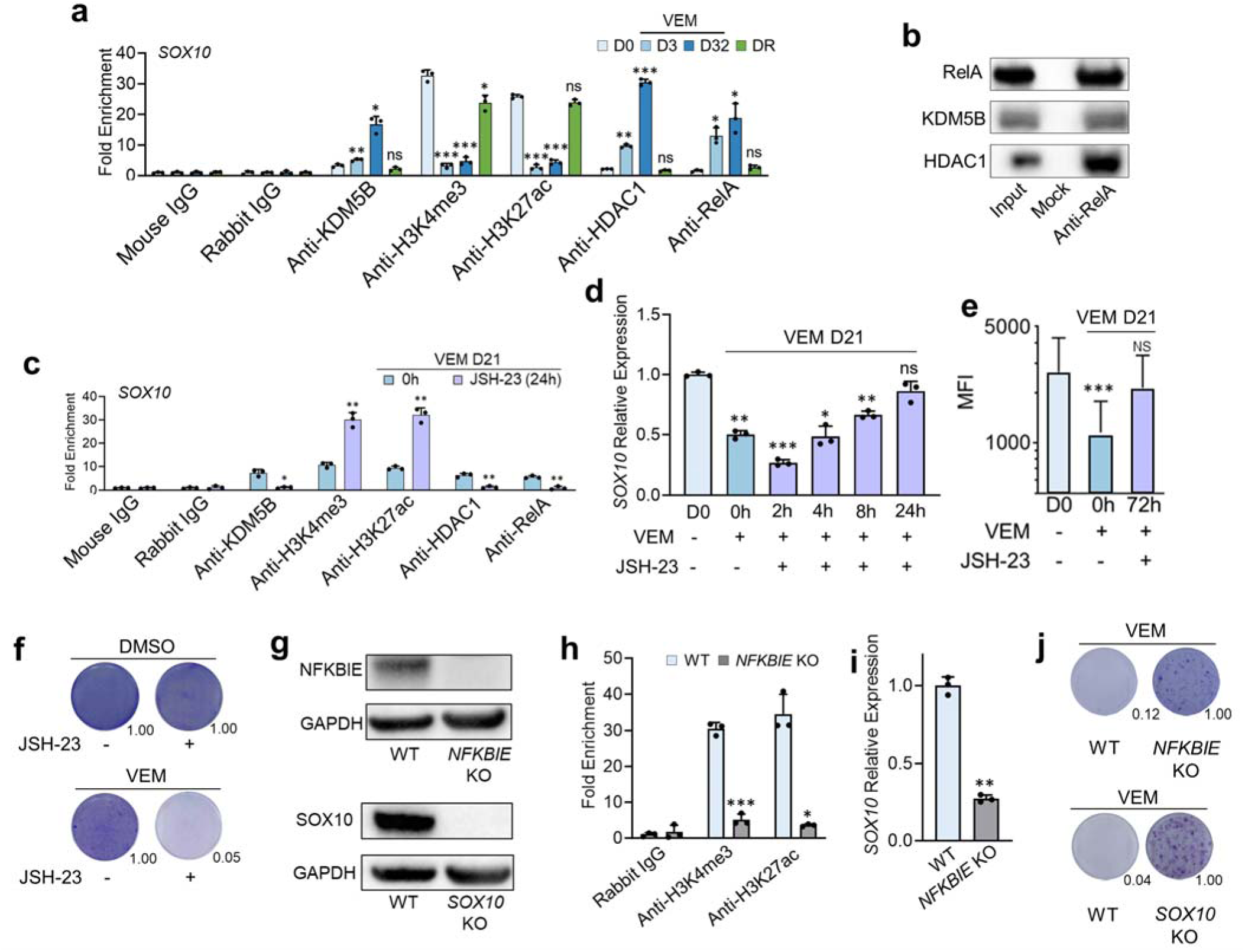
Mechanistic validation of RelA-driven chromatin remodeling underlying the dedifferentiation transition in M397 cells. **a.** ChIP-qPCR assessment of the binding profiles of RelA, KDM5B, HDAC1, H3K4me3, and H3K27ac at the promoter region of *SOX10* at selected time points across the dedifferentiation transition (mean ± SD, *p<0.05, **p<0.005, ***p<0.0005, NS: not significant, compared to D0, n=3). **b.** Co-immunoprecipitation of RelA with KDM5B and HDAC1, confirming the binding between RelA and the two histone modifiers. **c.** ChIP-qPCR assessment of the binding profiles of RelA, KDM5B, HDAC1, H3K4me3, and H3K27ac at the prompter region of *SOX10* before (0h) and after (24h) VEM+JSH23 treatment for M397 cells pretreated with VEM for 21 days (mean ± SD, ***p<0.0005 compared to respective control (0h), n=3). **d.** Recovery of *SOX10* gene expression levels in M397 cells pretreated with VEM for 21 days (D21), and then co-treated with VEM and JSH-23 for the indicated times, quantified by qPCR (mean ± SD, Welch ANOVA with correction for multiple comparisons using Dunnett T3, *p<0.05, **p<0.005, ns: not significant, compared to D0, n=3). **e.** SOX10 protein levels in M397 cells under the indicated treatment conditions assessed by flow cytometry. The reduced SOX10 protein expression after 21 days of VEM treatment was recovered after 72 hours VEM+JSH-23 treatment (mean ± SD, ***p<0.0005, compared to D0, NS: not significant). **f.** Clonogenic assays of M397 cells treated with VEM monotherapy or VEM+JSH-23 combination therapy for the same duration. Image quantification is shown in the lower right of each image. Toxicity of JSH-23 was assessed by comparing cells treated with JSH-23 or DMSO control for the same duration. **g.** Immunoblotting showing the successful KO of *SOX10* and *NFKBIE* in M397 cells. **h.** ChIP-qPCR assessment of H3K4me3 and H3K27ac levels at the promoter region of *SOX10* in *NFKBIE* KO M397 cells (mean ± SD, **p<0.005, ***p<0.0005 compared to wild-type (WT), n=3). **i.** Relative *SOX10* expression of *NFKBIE* KO M397 cells compared to wild type (WT) assessed by qPCR (mean ± SD, ***p<0.0005 compared to WT, n=3). **j.** Clonogenic assays of WT, *NFKBIE* KO, and *SOX10* KO M397 cells treated with 3 μM VEM for the same duration, with image quantification showing in the lower right.

To further validate this mechanism, we block RelA nuclear translocation using the inhibitor JSH-23^43^ in M397 cells with reduced *SOX10* expression levels after being treated with VEM for 21 days. Consistent with our hypothesis, this blockade reduced the RelA binding and its recruitment of the KDM5B and HDAC1, resulting in increased H3K4me3 and H3K27ac marks at the *SOX10* promoter (Fig. 5c). This led to the restoration of *SOX10* gene and protein expression shortly after JSH-23 treatment (Fig. 5d, e). These results support the notion of *SOX10* repression through RelA-mediated epigenetic silencing. Consistently, blocking RelA nuclear translocation with JSH-23 enhanced the therapeutic efficacy of BRAFi, resulting in more sustained growth inhibition (Fig. 5f). To further validate the role of RelA through orthogonal methods, we promoted RelA nuclear translocation by using CRISPR to knockout (KO) *NFKBIE*, thereby releasing the cytoplasmic retention of RelA (Fig. 5g). In line with our hypothesis, RelA translocation in *NFKBIE* KO cells reduced activation histone mark levels at the promoter of *SOX10* and decreased *SOX10* expression (Fig. 5h, i). Functionally, both *NFKBIE-*KO and *SOX10-*KO M397 cells developed drug tolerance to BRAFi VEM more rapidly than their wild-type counterparts (Fig. 5g, j). Collectively, these results reveal a novel mechanism where ROS-mediated RelA activation functions as an early-acting regulatory process that epigenetically suppresses the key melanocytic lineage TF SOX10, leading to melanoma cell state dedifferentiation towards BRAFi resistance.

### Generality of chromatin remodeling-mediated sequential transcriptional dynamics quantitatively reveals the epigenetic nature of cancer cell plasticity

To assess whether the sequential operation of M_early_ and M_late_ gene modules extends beyond M397, we analyzed temporal transcriptome data from several *BRAF*-mutant melanoma lines with varying degrees of plasticity, as collected in a previous study^44^. These lines exhibited phenotypic composition ranging from largely mesenchymal (e.g. M381) to in-between neural crest and melanocytic (e.g. M263), to predominantly melanocytic (e.g. M229). We projected the VEM-induced transcriptional dynamics of these lines onto the 2D space defined by M_early_ and M_late_. Upon BRAF inhibition, all lines exhibited the sequential operation of M_early_ and M_late_, resembling M397 (Fig. 6a), implicating mechanistic similarities in their drug-induced dedifferentiation trajectories.

**Fig. 6.**
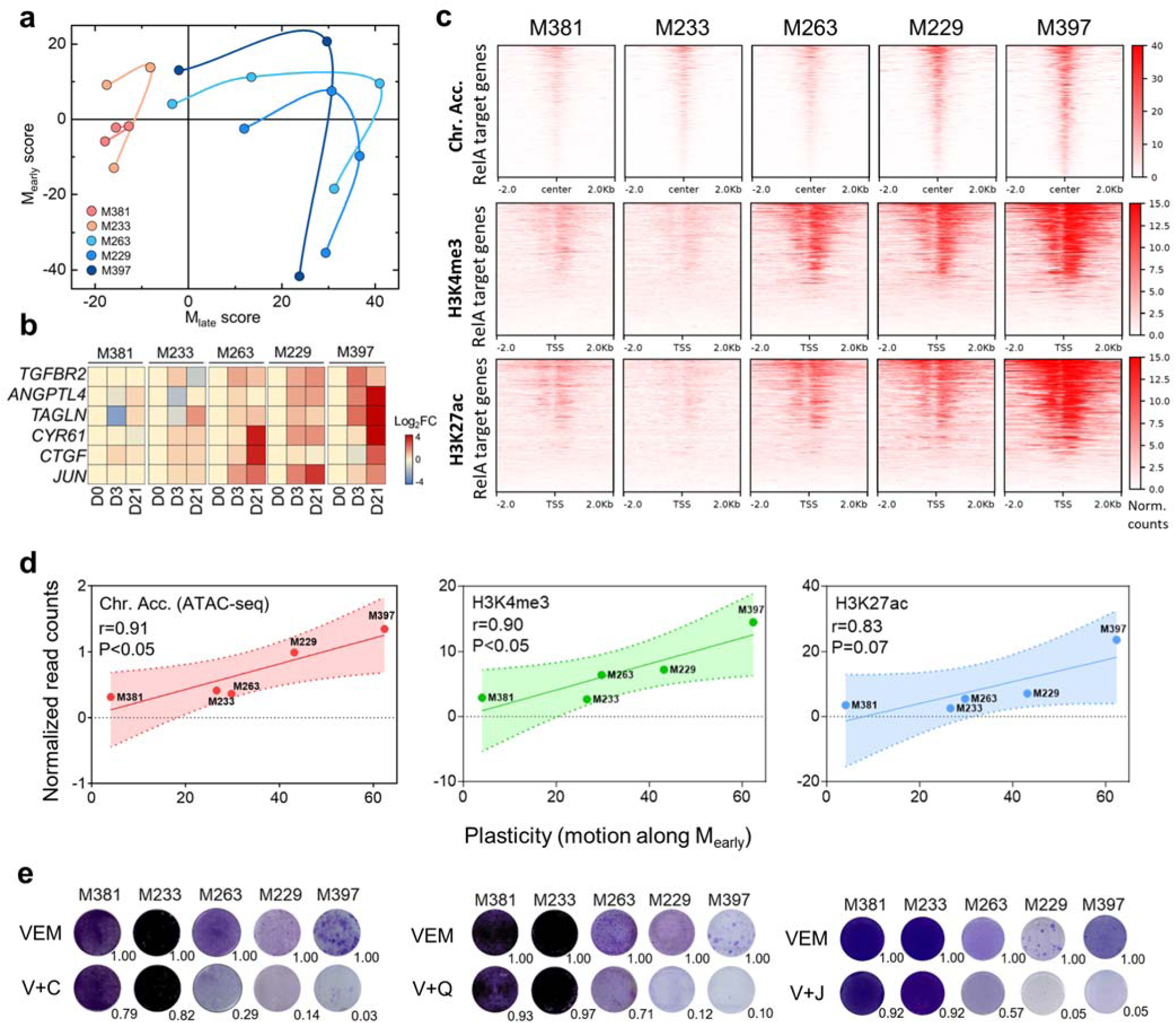
Generality of the sequential transcriptional dynamics and molecular underpinning of the phenotypic plasticity. a. Quantification of phenotypic plasticity upon BRAF inhibition across a panel of *BRAF*-mutant melanoma cell lines. Transcriptome data are projected to the 2D plane defined by M_early_ and M_late_, connected by smooth lines. Data points for each cell line represent day-0 (D0), D3, D21, D60 (for M229), and D90 (for M263) of BRAFi (VEM) treatment in counterclockwise order. b. Heatmap showing the Log2 fold changes (Log2FC) in expression levels of TGF-β receptor 2 and several bona fide TGF-β target genes at different time points of VEM treatment relative to untreated control (D0) across different melanoma cell lines. c. Heatmaps showing the normalized read counts of baseline chromatin accessibility (peak center ± 2kb), as well as H3K4me3 and H3K27ac levels at the promoter regions (TSS ± 2kb) across the 307 RelA target genes in different melanoma lines. d. Correlation between the average ATAC-seq, H3K4me3, and H3K27ac signals associated with the 307 RelA target genes and short-term plasticity (motion along M_early_) across different melanoma cell lines. Pearson correlation coefficients (r) and p-values are shown. The shaded regions denote 95% CI of the linear fitting. e. Clonogenic assays for VEM monotherapy (dosing varies across cell lines, see Methods) and combination therapies targeting the driver oncogene *BRAF* by VEM and RelA nuclear translocation by JSH-23 (V+J), histone modifiers KDM5B by CPI-455 (V+C), or HDAC1 by Quisinostat (V+Q) across different melanoma cell lines. Treatments were stopped upon clear cell regrowth in the VEM monotherapy groups. Image quantitation relative to BRAFi monotherapy treatment is shown in the lower right of each image. The cell lines are ordered from left to right with increasing plasticity.

However, different cell lines also displayed widely distinct amplitudes of motion within this 2D space and varying degrees of elevation in TGFβ pathway genes after 3 weeks of VEM treatment (Fig. 6a, b), reflecting substantial variations in their transcriptome plasticity. Notably, early-stage plasticity, evaluated by the drug-induced motion along M_early_, correlated with baseline chromatin accessibility and baseline levels of H3K4me3 and H3K27ac at the promoter regions across the 307 dedifferentiation-associated RelA target genes (Fig. 6c, d). Consistently, continued VEM exposure reduced the expression of most RelA target genes in highly plastic lines, such as M397 and M229, but had little impact on the least plastic line M381, which exhibited the least accessible chromatin at the promoters of these RelA target genes (Extended Data Fig. 6a). These positive correlations implicate a similar role of RelA-driven chromatin remodeling across these lines upon BRAF inhibition and suggest that the level of plasticity may be pre-encoded (epigenetically constrained) by the baseline chromatin permissiveness of these critical RelA target genes prior to treatment. This finding quantitatively reveals the epigenetic nature of cancer cell plasticity.

The phenotype plasticity of M397 cells upon BRAF inhibition is mediated by NF-κB/RelA-driven chromatin remodeling, which leads to the marked reduction in activation histone marks and chromatin accessibility, consequently altering the expression of the dedifferentiation-associated RelA target genes (Figs. 3d–f and 4a). This suggests that drug targeting the RelA nuclear translocation or histone modifiers to maintain the chromatin openness may arrest the BRAFi-induced dedifferentiation transition in the most plastic cell lines (e.g. M229, M397), but would have little effect on the less plastic cell lines (e.g. M381). Consistent with our prediction, for plastic melanoma cell lines that typically develop adaptive drug tolerance to VEM monotherapy within a few weeks of drug exposure, functional clonogenic assays revealed that drug combinations co-targeting RelA nuclear translocation or histone modifiers KDM5B and HDAC1, in conjunction of VEM, sufficiently inhibited the cell growth at doses without significant cytotoxicity when VEM monotherapy-treated persister cells began to regrow (Fig. 6e and Extended Data Fig. 6b)^45,46^. In contrast, these combinations provided little benefit over VEM monotherapy to cell lines exhibiting low baseline chromatin accessibility and minimal transcriptome plasticity, such as M381 and M233 (Fig. 6e and Extended Data Fig. 6b). These results highlight the potential utility of co-targeting the driver oncogene *BRAF* along with chromatin-remodeling machinery to treat those *BRAF*-mutant melanomas with permissive chromatin states poised to be remodeled upon BRAF inhibition.

### The pivotal role of oxidative stress-mediated NF-**κ**B/RelA activation in the transition towards DTP states in other cancer types

To explore the role of oxidative stress-mediated NF-κB/RelA activation in driving cell state changes towards DTP states in other cancer types, we studied the *EGFR*-mutant lung cancer cell line HCC827 and the *BRAF*-mutant colon cancer cell line HT-29. Previous reports have demonstrated that both lines can transition to DTP states after 9 days of sustained inhibition of driver oncogenes^4,47^. Specifically, HCC827 cells were treated with the EGFR inhibitor erlotinib, while HT-29 cells were treated with BRAFi dabrafenib and the EGFR antibody cetuximab (DAB+CET), and both lines reverted to their original untreated states upon drug removal^4,47^. We confirmed the generation of DTPs in both lines, with persister cell numbers constituting very small fractions of DMSO-treated controls (Fig. 7a). While HCC872 and HT-29 cells originate from epithelial lineages, distinct from the melanocytic lineage of M397 cells, both lines exhibited activation of the TGFβ pathway similar to M397 cells when transitioning to the DTP state. This was evidenced by elevated TGFβ pathway-related genes and increased phosphorylation of SMAD2 and c-Jun (Fig. 7b, c).

**Fig. 7.**
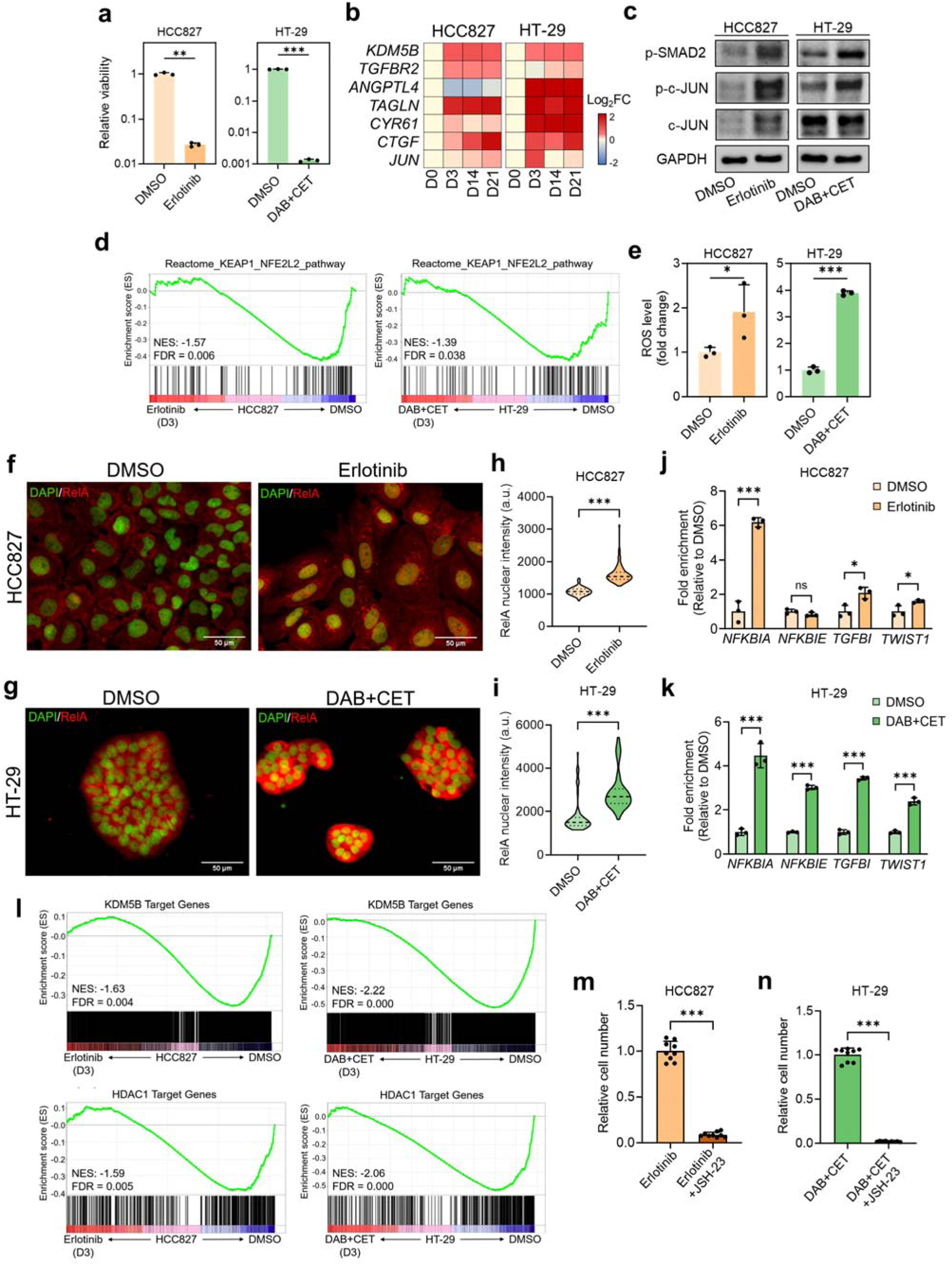
Oxidative stress-mediated NF-κB/RelA activation in the transition towards DTP states in other cancer types. **a.** Relative viability of DTPs compared to DMSO controls for HCC827 cells treated with erlotinib and HT-29 cells being treated with dabrafenib plus cetuximab (DAB+CET) for 9 days. Drugs were replenished every 3 days (mean ± SD, **p<0.005, ***p<0.0005, n=3). **b.** Heatmap showing the Log2 fold changes (Log_2_FC) in expression levels of KDM5B, TGF-β receptor 2, and several bona fide TGF-β target genes at different time points of drug treatment relative to control (D0) in HCC827 and HT-29 cells, with treatment conditions specified in (**a**). **c.** Immunoblotting showing the p-SMAD2 (Ser465/467), p-c-Jun (Ser63), and c-Jun levels in HCC827 and HT-29 cell lines after 9 days of drug treatment, with treatment conditions specified in (**a**). **d.** Gene set enrichment analysis showing the significant downregulation (negative enrichment) of NRF2-related pathway genes in HCC827 and HT-29 cells after 3 days of treatment with erlotinib or DAB+CET, respectively, compared to DMSO controls. NES: normalized enrichment score **e.** Change in cellular ROS levels in HCC827 and HT-29 cells upon 3 days of drug treatment relative to the DMSO control, with treatment conditions specified in (**a**) (mean ± SD, *p<0.05, ***p<0.0005, n=3). **f, g.** Representative fields of view of immunofluorescence staining showing increased RelA nuclear translocation in HCC827 (**f**) and HT-29 (**g**) cells upon 3 days of drug treatment compared to DMSO controls, with treatment conditions specified in (**a**). **h, i.** Quantitation of RelA nuclear fluorescence intensity in HCC827 (**h**) and HT-29 (**i**) cells after drug treatment relative to DMSO controls, with treatment conditions specified in (**a**) for (***p<0.0005, n=82 fields of view). **j, k.** ChIP-qPCR assessment of RelA binding at the promoter regions of several canonical RelA target genes in HCC827 (**j**) and HT-29 (**k**) cells after 3 days of drug treatment compared to DMSO controls, with treatment conditions specified in (**a**) (mean ± SD, *p<0.05, **p<0.005, ***p<0.0005, n=3). **l.** Gene set enrichment analysis showing the significant downregulation (negative enrichment) of KDM5B and HDAC1 target genes in HCC827 and HT-29 cells upon 3 days of drug treatment compared to DMSO controls, with treatment conditions specified in (**a**). NES: normalized enrichment score. **m.** Relative viability of HCC827 cells being treated by erlotinib monotherapy or in combination with RelA nuclear translocation inhibitor, JSH-23. Treatment was stopped when clear cell regrowth was observed in the erlotinib group, and cell numbers were assessed by nuclear staining and counting (mean ± SD, ***p<0.0005, n=9 fields of view). **n.** Relative viability of HT-29 cells being treated by DAB+CET or in combination with RelA nuclear translocation inhibitor, JSH-23. Treatment was stopped when clear cell regrowth was observed in the DAB+CET group, and cell numbers were assessed by nuclear staining and counting (mean ± SD, ***p<0.0005, n=9 fields of view).

In alignment with observations in M397 cells, we observed reduced expression of most genes critical to antioxidant defense and significant downregulation of the NRF2 pathway in HCC827 and HT-29 cells upon 3 days of drug treatment, highlighting the dysregulated antioxidant response in both cell lines (Fig. 7d and Extended Data Fig. 7a). Consequently, both lines showed elevated ROS levels, which triggered RelA nuclear translocation within 3 days of oncogene inhibition (Fig. 7e-i). Enhanced RelA nuclear signals were accompanied by increased binding of RelA to its canonical target genes, such as *NFKBIA* and *TWIST1*, in both lines (Fig. 7j, k)^48–50^, supporting RelA transcriptional activation during this early-stage of drug exposure. Notably, similar to M397 cells, we also observed consistently elevated KDM5B expression levels during drug treatment in both HCC827 and HT-29 cells (Fig. 7b). Enrichment analysis further demonstrated significant downregulation of KDM5B and HDAC1 target genes after 3 days of drug treatment compared to DMSO control (Fig. 7l), suggesting the potential involvement of these two histone modifiers in epigenetically repressing their target gene expression at the onset of the transition.

Consequently, similar to the findings in *BRAF*-mutant melanoma lines, blocking RelA nuclear translocation using JSH-23 at doses without significant cytotoxicity in combination with erlotinib or DAB+CET resulted in a sustained cell growth inhibition over an extended period, effectively suppressing the establishment and regrowth of DTPs (Fig. 7m, n and Extended Data Fig. 7b, c). Collectively, these results underscore the commonality of oncogene inhibition-induced transitions towards DTP states across different cancer types and highlight the pivotal role of oxidative-stress mediated NF-κB/RelA activation in promoting such transitions.

## Discussion

Sequential transcriptional waves mediated by epigenetic reprogramming have been observed in embryonic development and T cell differentiation^1–3^. Here we have demonstrated that cancer cells can exploit similar transcriptional dynamics but in a reverse way (dedifferentiation process) to escape oncogene inhibition in a melanoma model of BRAFi resistance. Our information theoretic analysis revealed the hysteretic nature of the cell-state transition, highlighting the sequential operation of two tightly-coupled transcriptional waves. The early-acting gene module, associated with chromatin remodeling, exerts dominant epigenetic control over the full cycle of cell state changes. This picture guided the identification of a mechanistic regulatory network underpinning the dedifferentiation transition. Here, oxidative stress-mediated NF-κB/RelA transcriptional activation recruits histone erasers KDM5B and HDAC1 as co-factors to epigenetically repress a group of RelA target genes associated with neural crest differentiation. This leads to the reduced expression of critical melanocytic lineage TFs, particularly *SOX10* and *MITF*, the upregulation of TGFβ signaling, and the ultimate transition towards drug tolerance^15,18,51^. The hysteresis observed in the drug-induced cellular response, while not previously documented as a behavior of phenotype plasticity in cancer, may indeed be common, given that chromatin serves as the essential medium through which TFs and other coregulators alter gene activity to promote cell-state changes^52,53^.

While cancer cell plasticity has been investigated in various model systems, it has mostly been described qualitatively. Here, our identification of two gene modules provides a quantitative measure to gauge the levels of plasticity in different *BRAF*-mutant melanoma lines at whole transcriptome scale. The projection of their BRAFi-induced transcriptome dynamics revealed that, although the sequential order of operation of the two modules is conserved across cell lines, the magnitudes of motion, reflecting the levels of plasticity, vary significantly. Surprisingly, the cell plasticity is strongly correlated with chromatin accessibility and activation histone marks of critical RelA target genes prior to drug treatment, implying that cellular plasticity may be epigenetically encoded prior to drug exposure. Cells with more permissive baseline chromatin states are poised with a ‘hair-trigger’ response to drug challenge, with the early-acting gene module setting in motion epigenetic and transcriptional programs for longer-term changes that ultimately result in the dedifferentiated drug-resistant phenotype.

How these bulk-level observations extend to single-cell resolved studies remains unclear and represents an exciting direction for future investigation. Single cells may exhibit a distribution of permissive chromatin states, with the most permissive being the most susceptible to dedifferentiation towards drug-tolerance. New tools for the epigenetic analysis of single cells *in vitro* and in spatially resolved tissues offer opportunities for resolving this heterogeneity^54–56^. These findings also support the use of epigenetic drugs to potentiate oncogene inhibition, effectively expanding the drug arsenal against cancer progression^57^. Indeed, targeting histone-modifying enzymes KDM5B and HDAC1 in combination with BRAFi led to sustained growth inhibition in plastic melanoma cells *in vitro*. While *in vitro* models may not fully recapitulate the cellular behavior *in vivo*, the presence of the dedifferentiation transition in melanoma mouse xenografts and patient samples resistant to BRAFi treatments suggests consistency, warranting further exploration (Fig. 1d–g and Extended Data Fig. 2).

NF-κB signaling has long been recognized as a crucial link between inflammatory microenvironment and cancer progression^58^. Our work demonstrates a novel alternative role of NF-κB activation, connecting cellular oxidative stress with cancer progression by driving adaptive cell-state transitions towards DTPs across different cancer types. Activation of NF-κB/RelA through either ROS-induced direct phosphorylation of iκB family proteins or ROS-mediated inactivation of serine/threonine phosphatases offers enhanced protection from oxidative-stress-induced cell death. This represents an essential mechanism that cells utilize to respond to an adverse environment^37,38,42,59,60^. We observed that oncogene inhibition disrupted the redox balance and triggered dysregulated antioxidant defense, leading to the ROS accumulation within days of drug exposure across these cancer models of adaptive resistance. Consequently, we found consistent activation of the built-in protection mechanism, NF-κB/RelA signaling, across these cancer models, as evidenced by the elevated RelA nuclear translocation and enhanced binding to the promoters of its canonical target genes.

In M397 melanoma cells, RelA activation recruited histone modifiers to remodel the chromatin of lineage-associated TFs at the early stage of drug treatment, promoting cell-state dedifferentiation towards the DTP state with BRAFi resistance. While HCC827 and HT-29 cells have different tissue origins and lineage specifications from melanoma, the elevated KDM5B expression levels and highly enriched changes of KDM5B and HDAC1 target genes upon 3 days of drug treatment suggested the potential involvement of the same histone modifiers in initiating adaptive transitions in these models. Furthermore, the ROS-mediated RelA nuclear translocation and the inhibition of this translocation early on, which could prevent the emergence of DTPs and induce a sustained growth inhibition, suggest a common and indispensable role for NF-κB/RelA signaling in fostering the transitions toward DTP states across these models.

However, further work is required to determine whether RelA interacts with these histone modifiers in the same way as observed in M397 cells to regulate critical downstream lineage TFs, or whether other potential epigenetic mechanisms are involved in this process. Resolving the specific downstream TFs driving adaptive cell state changes in HCC827 and HT-29 is beyond the scope of this study. However, consistent with plastic melanoma lines, these models also showed the activation of TGFβ signaling upon transitioning to persister states. This is another commonality that could be therapeutically exploited to eliminate the DTPs.

Collectively, our results highlight the pivotal role of the NF-κB/RelA axis in driving the adaptive transition towards DTPs across multiple cancer types and lineage specifications. These insights would have been difficult to uncover without high-resolution temporal multi-omic characterization and a systems-level computational framework to model the transition dynamics and distill the critical regulators.

## Methods

### Cell lines and drug treatment

M-series patient-derived cell lines used in this study were generated under UCLA institutional review board approval No. 11–003254. Cells were cultured in a humidified incubator at 37°C with 5% CO2 using RPMI 1640 with L-glutamine (Life Technologies), supplemented with 10% fetal bovine serum (Omega) and 0.2% antibiotics (MycoZap™ Plus-CL, Lonza). These cell lines were periodically tested for mycoplasma contamination and authenticated using the GenePrint 10 System (Promega) to ensure consistency with early passages. BRAF inhibitor (vemurafenib, Selleckchem S1267), KDM5B inhibitor (CPI-455, Selleckchem S8287), HDAC inhibitor (quisinostat, Selleckchem S1096), and RelA translocation inhibitor (JSH-23, Selleckchem S7351) were dissolved in DMSO at designated concentrations before application to the cell culture media. Cells were plated in 10 cm tissue culture dishes at optimal confluency and treated with the respective drugs for the designated durations. M397 cells were exposed to 3 µM vemurafenib for 59 days or for 29 days followed by a drug washout period and culture in drug-free medium for an additional 35 days. Vemurafenib was replenished three times per week. HCC827 (CRL-2868) and HT-29 (HTB-38) were purchased from ATCC. Cells were cultured in a water-saturated incubator at 37 °C with 5% CO_2_. HCC827 (CRL-2868) and HT-29 (HTB-38) cell lines were obtained from ATCC. HCC827 cells were cultured in ATCC-formulated RPMI-1640 Medium (ATCC 30-2001) and treated with 2 μM of the EGFR inhibitor erlotinib (Selleckchem S1023) for 3 weeks. HT-29 cells were cultured in ATCC-formulated McCoy’s 5a Medium Modified (ATCC 30-2007) and treated with 50 μg/mL of the anti-EGFR antibody cetuximab (Selleckchem A2000) in combination with 1 μM of the BRAF inhibitor dabrafenib (MedChemExpress HY-14660) for 3 weeks. Erlotinib and dabrafenib were replenished every 3 days, while cetuximab was replenished every 5 days. The drug dosages and treatment conditions used for HCC827 and HT-29 cells to generate drug-tolerant persisters (DTPs) were based on previous studies^4,47^. Treated cells were harvested at selected time points for the time-series transcriptome analyses by RNA sequencing.

### Melanoma orthotopic mouse xenograft models

Mice were housed in a specific pathogen-free (SPF) animal facility at the University of Washington, maintained at 22°C ± 1°C, on a reverse 12-hour light/dark cycle. Food and water were provided *ad libitum*. The mice were kept under animal bio-safety level 2 (ABSL-2) conditions and handled according to protocols approved by the Institutional Animal Care and Use Committee (IACUC) of the University of Washington, in compliance with international guidelines. To generate xenograft mouse tumor models, 6-week-old NOD.*Cg-Prkdc^scid^Il2rg^tm1Wjl^*/SzJ (NSG) female mice were purchased from the Jackson Laboratory (Cat#: 005557). NSG mice were inoculated subcutaneously into the right flank with M397 cells (8 × 10^5^ cells per mouse) in Matrigel mixture (Corning cat#: 354234). Once the average tumor volume reached approximately 200 mm³, the mice were randomized into four groups (n = 3 per group): control, 50 mg/kg vemurafenib, 100 mg/kg vemurafenib, and 200 mg/kg vemurafenib. Vemurafenib (LC Laboratories, Cat#: V-2800) was prepared as a 5% DMSO solution in 1% methyl cellulose and administered once daily by oral gavage (p.o.). Tumor dimensions were measured every three days using calipers. Tumor volume (V) was calculated using the formula: V (mm³) = Length (mm) × Width (mm²) / 2. Mice with tumor volumes exceeding 1500 mm³ were euthanized, and tumors were excised. The excised tumors were fixed in 10% neutral buffered formalin and embedded in paraffin. Formalin-fixed, paraffin-embedded (FFPE) tumor samples were sent to the Translational Pathology Core Laboratory (TPCL) at the University of California, Los Angeles for multiplex immunohistochemistry (mIHC) staining.

### Melanoma patient samples

Pretreatment melanoma samples were obtained from surplus biopsies stored in the melanoma biobank at the Peking University Cancer Hospital and Institute (Beijing, China). Patient #1 was treated with vemurafenib, while Patient #2 received a combination of dabrafenib and trametinib. Both patients initially exhibited a partial response (PR) to these BRAF (or MAPK) inhibitors. Secondary biopsies were collected when the patients’ disease progressed to a state of progressive disease (PD). Informed consent was obtained from both patients for the use of their biopsy materials in scientific research. All research activities were conducted following the guidelines and protocols approved by the institutional ethics review committee, adhering to local laws for research on human-derived tissues. The gender of the patients was as follows: Patient #1, female; Patient #2, female.

### RNA-seq library preparation

Total RNA from M-series melanoma cells was extracted using the RNeasy Mini Kit (Qiagen). For unstranded libraries, transcriptome libraries were constructed with the TruSeq RNA Sample Preparation Kit V2 (Illumina, San Diego, USA) with minor modifications. Specifically, 250 ng of total RNA from each sample was used to construct cDNA libraries, which were then sequenced on the Illumina HiSeq 2500 platform using single-end 51 bp reads, following the manufacturer’s recommendations. For stranded libraries, RNA sequencing libraries were prepared using the Kapa RNA mRNA HyperPrep Kit (Kapa Biosystems) according to the manufacturer’s protocol. Briefly, 100 ng of total RNA from each sample was used for polyA RNA enrichment with magnetic oligo-dT beads. The enriched mRNA underwent fragmentation using heat and magnesium, and first-strand cDNA was synthesized using random priming. The combined second-strand cDNA synthesis with dUTP and A-tailing reaction generated double-stranded cDNA with dAMP at the 3’ ends. Barcoded adaptors (Illumina) were then ligated to the double-stranded cDNA fragments. For HCC827 and HT-29 cells, total RNA was extracted from cell pellets using the AllPrep DNA/RNA Kit (Qiagen). mRNA libraries were constructed using the TruSeq RNA Sample Preparation Kit V2 (Illumina, San Diego, USA) and sequenced on the Illumina HiSeq 2500 platform with paired-end 150 bp reads, following the manufacturer’s recommendations. Sequencing was conducted by Novogene.

### ATAC-seq library preparation

ATAC-seq data was generated following a previously published protocol for cell lysis, tagmentation, and DNA purification^61^. The Tn5-treated DNA was initially amplified using a 5-cycle PCR in 50 µl reaction volumes. After this initial amplification, 5 µl of the partially amplified library was used to perform qPCR to determine the number of additional PCR cycles needed. Based on the qPCR results, an additional 4-5 PCR cycles were performed on the remaining 45 µl of each partially amplified product. Final PCR cleanup was conducted using AmpureXP beads purification. The resulting libraries were validated using the Agilent Bioanalyzer DNA High Sensitivity Kit and quantified by qPCR. ATAC-seq library templates were prepared using the Illumina HiSeq PE Cluster V4 Kit.

### ChIP-seq library preparation and ChIP-qPCR

Chromatin immunoprecipitation (ChIP) for H3K4me3, H3K27ac, NF-κB p65/RelA, KDM5B, and HDAC1 was performed using the Magna ChIP A/G Chromatin Immunoprecipitation Kit (Millipore Cat# 17-10085). Briefly, cells were cultured to approximately 80% confluency in a petri dish containing 10 mL of growth media. Cells were then fixed with 1% formaldehyde by adding 275 µl of 37% formaldehyde for 10 minutes at room temperature to cross-link protein-DNA complexes. The unreacted formaldehyde was quenched by adding glycine to a final concentration of 0.125 M and gently swirling the dish to mix. The nuclear pellet was isolated using Cell Lysis Buffer and resuspended in 500 µl SDS Lysis Buffer containing 1X Protease Inhibitor Cocktail II. Sonication was performed for 4 minutes (10 seconds on, 30 seconds off, 10% strength) using a Bioruptor to yield DNA fragments of 0.2-1.0 kb in length. The lysates were cleared by centrifugation (12,000g for 10 minutes at 4°C) and diluted tenfold in ChIP Dilution Buffer to reduce the SDS concentration. After reserving 10% of the sample as input, 500 µl of the supernatant was incubated overnight at 4°C with specific antibodies (anti-H3K4me3, Millipore Cat# 07-473; anti-H3K27ac, Millipore Cat# 17-683; anti-RelA, Millipore Cat# 17-10060; anti-JARID1B/KDM5B, Bethyl Lab, Cat# A301-813A; anti-HDAC1, Millipore Cat# 17-608) and 20 µL of fully resuspended protein A/G magnetic beads (Millipore Cat# 16-663). Washing, elution, reverse cross-linking, and purification steps were performed according to the manufacturer’s instructions. The eluted DNA was quantified using the Qubit dsDNA HS Assay Kit (Invitrogen Cat# Q32851) and used for subsequent ChIP-qPCR or ChIP-seq library preparation.

ChIP-seq libraries were prepared using the Kapa DNA HyperPrep Kit (Kapa, Cat# KK 8700) following the manufacturer’s protocol. Briefly, 5-10 ng of immunoprecipitated DNA underwent end-repair, A-tailing, and adaptor ligation. Subsequently, a 10-cycle PCR was performed to produce the final sequencing library. The libraries were validated using the Agilent Bioanalyzer DNA High Sensitivity Kit and quantified with Qubit. For ChIP-qPCR, the SsoAdvanced Universal SYBR Green Supermix (Bio-Rad, Cat# 1725271) was used on a CFX96 Real-Time PCR Detection System. In each PCR/qPCR reaction, 2 µl of eluted DNA was added. The primer sequences (IDT) used for qPCR were as follows: *SOX10* FW: 5’-CCCGACTCATGCTGCCAA-3’, *SOX10* REV: 5’-TCTGGCCTGGGTAGAAGGG-3’; *NFKBIE* FW: 5’-TGTGTGCATTGGGGCTTTTT-3’, *NFKBIE* REV: 5’-GAACTACTGGCCTCTGACCC-3’, NFKBIA REV: 5’-AGCTCGTCCGCGCCATGTTCCAG-3’; TGFBI FW: 5’-TGCGGAAGGTCAGGTAGTC-3’, TGFBI REV:5’-GCAAGGTTTTTAGTTCGGGC-3’; TWIST1 FW: 5’-TGGAGACTGCTCTCCCTAGC-3’, TWIST1 REV: 5’-TTCCCAGTCCAC CTCGATTTCC-3’.

### Sequencing ChIP-seq and ATAC-seq library

Sequencing runs were performed on the Illumina HiSeq 2500 platform in single-read mode with 51 cycles for Read 1 and 7 cycles for the index read, using SBS V4 Kits. For sequencing ChIP-seq biological replicates, the Illumina NovaSeq 6000 was utilized with the S4 Kit v1.5, in paired-end mode with 2 × 101 cycles. For ATAC-seq, sequencing runs were carried out in paired-end mode with 101 cycles on the Illumina HiSeq 2500 platform using HiSeq SBS V4 Kits. Real-time analysis (RTA) software version 2.2.38 was used for image analysis, and base calling was performed using bcl2fastq. Depending on the specific sequencing run, either bcl2fastq version 1.8.4 or 2.18 was used for samples processed at the City of Hope.

### Cell Viability Assay

For the cell viability assay, M397 cells were seeded into each well of a 96-well plate at an appropriate confluency and treated with the indicated drug concentrations for 72 hours. Cell viability was measured using the ATP-based CellTiter-Glo Luminescent Assay (Promega, Cat# G7570), which quantifies cell number and allows for the construction of dose-response curves. IC50 values were calculated from at least three biological replicates. HCC827 and HT-29 cells were seeded onto 10 cm petri dishes in biological triplicate at an appropriate confluency. HCC827 cells were treated with DMSO or 2 μM erlotinib, and HT-29 cells were treated with DMSO or a combination of 50 μg/mL cetuximab plus 1 μM dabrafenib, for 9 days. Relative viabilities were assessed through cell counting. Erlotinib and dabrafenib were replenished every 3 days, while cetuximab was replenished every 5 days.

### Cell cycle assay

For cell cycle analysis, 500,000 cells were plated and treated with EdU. After treatment, the cells were washed with PBS and fixed. EdU detection was performed using the Click-iT EdU Alexa Fluor 488 Flow Cytometry Assay Kit (Thermo Fisher Scientific, Cat# C10420) according to the manufacturer’s protocol. DNA content was visualized using SYTOX AADvanced (Thermo Fisher Scientific, Cat# S10274). Gates were determined using an unstained control. All experiments were performed with at least two biological replicates.

### Fluorescence imaging of cell lines

MITF and F-actin fluorescent micrographs of M397 cells were captured using a Nikon C2plus confocal microscope (Ti) equipped with a Plan Apo λ 20× objective (Nikon Inc., Melville, NY). The microscope was controlled by NIS Elements AR software (version 4.51.00). Imaging settings were as follows: 30 μm pinhole, 12-bit acquisition, 25-30 PMT gain, and laser power of 0.7% (405 nm), 1.0% (488 nm), and 0.4% (640 nm). Cells were adhered to gelatin-coated glass surfaces in 96-well glass-bottom plates (Greiner Sensoplate Plus, Cat# 655892). To prepare the surface, 100 µL of 0.1% gelatin solution was added to each well and incubated at room temperature for 10 minutes. After incubation, the gelatin solution was removed, and the wells were air-dried for at least 15 minutes. Approximately 10,000 cells were seeded per well in 100 µL of culture media and grown to ∼70% confluency. To fix the cells, an equal volume of 4% PFA solution was gently added to each well. The cells were fixed for 20 minutes at room temperature, then washed twice with wash buffer (0.1% BSA in PBS). Cells were then blocked and permeabilized in blocking buffer (10% normal donkey serum, 0.3% Triton X-100 in PBS) for 45 minutes at room temperature. After removing the blocking buffer, cells were incubated with mouse anti-MITF primary antibodies (Thermo Fisher Scientific, Cat# MA5-14154) diluted to 5 μg/mL in antibody diluent (1% BSA, 1% normal donkey serum, 0.3% Triton X-100 in PBS) for 4 hours at room temperature. Following two washes with wash buffer, cells were incubated with donkey anti-mouse IgG, Alexa Fluor 647 secondary antibody (Thermo Fisher Scientific, Cat# A31571, RRID: AB_162542) diluted to 4 μg/mL in antibody diluent for 1 hour at room temperature. After two more washes with wash buffer, cells were counterstained with Alexa Fluor 488 Phalloidin (Thermo Fisher Scientific, Cat# A12379) for 20 minutes at room temperature, following the manufacturer’s instructions. After additional washes with wash buffer, cells were further counterstained with 4’,6-Diamidino-2-Phenylindole (DAPI) (Thermo Fisher Scientific, Cat# D1306) diluted to 1 μg/mL in PBS for 5 minutes. Finally, after two PBS washes, the wells were filled with 78% glycerol for imaging.

The RelA nuclear translocation assay was conducted using fluorescence imaging with an NF-κB p65 (RelA) antibody and the ImageXpress XLS high-content fluorescence system (Molecular Devices). Cells were initially fixed with 4% paraformaldehyde for 15 minutes and then permeabilized with 0.1% Triton-100 for 15 minutes at room temperature. After three washes with PBS, the cells were blocked with 1% BSA in PBS for 1 hour at room temperature. Subsequently, cells were incubated with a 1:200 dilution of RelA primary antibody (Millipore #17-10060) in a solution of 0.1% BSA in PBS at 4°C overnight. The next day, cells were stained with Cy5 AffiniPure Donkey Anti-Mouse IgG (H+L) secondary antibody (Jackson ImmunoResearch Inc., Cat# 715-175-150) for 1 hour at room temperature. This was followed by another blocking step with 1% BSA in PBS for 1 hour at room temperature. Finally, the cells were stained with 1 μg/mL DAPI in PBS for 20 minutes at room temperature. After two washes with PBS, the cells were imaged using the ImageXpress system. The RelA signals in the cellular nuclei were quantified using nuclear masks generated by DAPI fluorescence signals, processed with MetaXpress software (Molecular Devices).

### Immunoblotting

Histone proteins were extracted using the Histone Extraction Kit (Abcam, Cat# ab113476). The following antibodies were used: H3K4me3 (Millipore, Cat# 07-473), H3K27ac (Millipore, Cat# 17-683), NFKBIE (Sigma-Aldrich, Cat# HPA005941), H3 (Abcam, Cat# ab18521), SOX10 (Cell Signaling Technology, Cat# 89356), GADPH (Cell Signaling Technology, Cat# 5174), phospho-IkappaB-ε (Sigma, Cat# SAB4503766), Phospho-SMAD2 (Cell Signaling Technology, Cat# S3108S), p-c-JUN (Santa Cruz Biotechnology, Cat# sc-822), and c-Jun (Santa Cruz Biotechnology, Cat# sc-74543). The Invitrogen precast gel system NuPAGE was used for SDS-PAGE with 4–12% Bis-Tris gels. Samples were loaded onto the gels, and after electrophoresis, the proteins were transferred to membranes. The membranes were blocked in a 5% BSA solution with TBS + 0.1% Tween-20 (TBST) for at least 1 hour at room temperature. Subsequently, the membranes were incubated overnight at 4°C with primary antibodies diluted in 5% BSA with TBST. The next day, the membranes were washed three times for 5 minutes each in TBST and then incubated with a suitable HRP-coupled secondary antibody for 1 hour at room temperature. After three additional washes, the proteins were visualized using SuperSignal™ West Pico PLUS Chemiluminescent Substrate (Thermo Fisher Scientific, Cat# 34577) and detected with the ChemiDoc™ XRS+ System.

### RT-qPCR

For quantitative reverse transcription-polymerase chain reaction (qRT-PCR), total RNA was extracted using the TRIzol Plus RNA Purification Kit (Thermo Fisher Scientific, Cat# 12183555) and reverse-transcribed into cDNA. Real-time PCR was performed with gene-specific primers on the two-color real-time PCR detection system (Bio-Rad) using the SsoAdvanced Universal SYBR Green Supermix (Bio-Rad, Cat# 1725272) to measure relative expression levels. The sequences of the primers (IDT) used were: SOX10 FW: 5’-CCTGAGGTGGGCAAGGAAC-3’; SOX10 REV: 5’-AGGACCCTATTATGGCCACTCG-3’; NFKBIE FW: 5’-CCCGGTTGGTCCAGATGTAC-3’, NFKBIE REV: 5’-GGGATCCCAAGAGCTTGGTG-3’.

### Co-IP and protein detection

For cell lysis, cells were cultured to approximately 80% confluency in a petri dish containing 10 mL of growth media and washed three times with ice-cold PBS. The cells were then collected with a scraper in 1 mL ice-cold PBS supplemented with 1X protease inhibitor cocktail (Cell Signaling Technology, Cat# 5871) and centrifuged. The cell pellets were resuspended in a cell lysis buffer containing 50 mM Tris-HCl (pH 7.5), 250 mM NaCl, 1 mM EDTA, 0.5% Triton X-100, 10% glycerol, and 1X protease inhibitor cocktail. The resuspended cell pellets were incubated at 4°C for 30 minutes and sonicated in an ice-water bath three times with 5-second pulses each. The cell lysates were then cleared by centrifugation at 10,000 × g for 10 minutes at 4°C. Protein concentration was quantified using the Qubit Protein Assay Kit (Invitrogen, Cat# Q33211).

For cross-linking the antibody to magnetic beads, 20 μl of magnetic protein A/G beads (Millipore, Cat# 16-663) were washed twice with cell lysis buffer and resuspended in 100 μl of cell lysis buffer without glycerol. Five micrograms of anti-NF-κB p65 (RelA) antibody (Millipore, Cat# 17-10060) were coupled to the magnetic protein A/G beads by incubating overnight at 4°C on a rotator. The RelA antibody-coupled protein A/G beads were washed three times in 200 µL of Conjugation Buffer (20 mM Sodium Phosphate, 0.15 M NaCl, pH 7.5). The beads were then resuspended in 250 µL of 5 mM BS3 (Thermo Fisher Scientific, Cat# 21585) with conjugation buffer and incubated at room temperature for 30 minutes with rotation. The cross-linking reaction was quenched by adding 12.5 μl of 1 M Tris-HCl (pH 7.5) and incubated at room temperature for 15 minutes with rotation. The RelA antibody-conjugated protein A/G beads were washed three times with cell lysis buffer.

For the co-immunoprecipitation (Co-IP) experiment, 200 μL of pre-cleared cell lysates were added to RelA antibody-conjugated protein A/G beads and incubated overnight at 4°C with rotation. The beads were then washed five times with 500 μL of cell lysis buffer without glycerol. The pellet beads were collected using a magnetic stand and resuspended in 65 μL of SDS buffer (50 mM Tris-HCl pH 6.8, 2% SDS, 10% glycerol, 1% β-mercaptoethanol). For immunoblotting, the eluted samples were boiled for 10 minutes at 95°C. Then, 20 μL of the boiled elutes were electrophoresed on 10% Mini-PROTEAN TGX Precast Gels with SDS running buffer. The proteins were transferred onto PVDF membranes using the Bio-Rad Wet Blotting Systems. The membranes were blocked with 5% non-fat dried milk (Bio-Rad) dissolved in PBS for 1 hour at room temperature and incubated overnight at 4°C with the following primary antibodies: JARID1B/KDM5B (Bethyl Lab, Cat# A301-813A), NF-κB p65/RelA (Millipore, Cat# 17-10060), and HDAC1 (Millipore, Cat# 17-608). After incubation, the membranes were treated with secondary goat anti-mouse/rabbit antibodies coupled with HRP (Thermo Fisher Scientific, Cat# 32430) and visualized using the ChemiDoc XRS+ Imaging System.

### CRISPR engineering of cell lines

For CRISPR engineering of cell lines, LentiCRISPR v2 plasmids targeting the coding sequences of *SOX10* or *NFKBIE*, along with a control LentiCRISPR v2 plasmid, were purchased from GenScript. Lentiviruses were produced in HEK-293T cells by transient transfection with the LentiCRISPR v2 plasmid and packaging vectors psPAX2 and pMD2.G, following previously described methods^62^. The virus was collected 48 hours post-transfection, filtered through a 0.45 µm syringe filter, and used to infect M397 cells. Infection was performed by spin-infection with viral supernatant supplemented with 10 µg/mL polybrene at 2,500 rpm and 30°C for 90 minutes. Transduced cells were selected using puromycin starting 3 days post-transduction. Genome editing at the respective loci was examined using a surveyor assay, performed according to the manufacturer’s instructions (Integrated DNA Technologies)^63^.

### Drug efficacy assays

For drug efficacy assays, melanoma cells were plated onto six-well plates with fresh media at an optimal confluence. In the clonogenic assays shown in Fig. 5f, M397 cells were treated with either 3 µM vemurafenib monotherapy or a combination of 3 µM vemurafenib and 25 µM JSH-23 for the same duration. The toxicity of JSH-23 was assessed by treating M397 cells with 25 µM JSH-23 or a DMSO control for the same duration. In the clonogenic assays shown in Fig. 5j, wild-type (WT), *NFKBIE* knockout (KO), and *SOX10* KO M397 cells were treated with 3 µM vemurafenib for the same duration. For the assays in Fig. 6e, each melanoma line was treated with vemurafenib at a concentration determined based on their IC50 value, as specified in previous work^5^, or in combination with 25 µM CPI-455, 20 nM Quisinostat, or 25 µM JSH-23 for the same duration. Treatments were stopped and followed by staining when vemurafenib-treated cells displayed clear regrowth. For toxicity assessments (Extended Data Fig. 6b and Fig. 7b), cells were treated with 25 µM CPI-455, 20 nM Quisinostat, or 25 µM JSH-23, compared to a DMSO control for the same duration. Media containing drugs or DMSO were replenished three times per week. At the time of staining, colonies were fixed with 4% paraformaldehyde and stained with 0.05% crystal violet solution. Image quantitation was performed using ImageJ.

In experiments shown in Fig. 7m, n, and Extended Data Fig. 7c, HCC827 and HT-29 cells were plated onto six-well plates with fresh media at an optimal confluence. HCC827 cells were treated with 2 μM erlotinib or a combination of 2 μM erlotinib and 25 µM JSH-23 for the same duration. HT-29 cells were treated with 50 μg/mL cetuximab and 1 μM dabrafenib, or a combination of these drugs with 25 µM JSH-23 for the same duration. Treatments were stopped when erlotinib-treated HCC827 cells or cetuximab plus dabrafenib-treated HT-29 cells displayed clear regrowth. The cells were then stained with Hoechst 33342 Solution (BD Bioscience, Cat# 561908) at a 1:100 dilution and imaged using a fluorescence microscope (Nikon ECLIPSE Ti Series) with nine randomly selected fields of view for each condition. Cell counting was performed using ImageJ based on the Hoechst nuclear staining.

### Protein phosphatase Assay

M397 cells were treated with 3 μM vemurafenib for 3 days and subsequently collected. Proteins were extracted using RIPA buffer (Cell Signaling Technology, Cat# 9806S), purified using spin columns from the Serine/Threonine Phosphatase Assay Kit (Promega, Cat# V2460) according to the manufacturer’s protocol, and quantified using the protein BCA assay. The assay determines the amount of free phosphate generated in a reaction by measuring the absorbance of a molybdate:malachite green:phosphate complex. Phosphate standards were generated by diluting 1 mM phosphate standard with the supplied phosphate-free water. Wells containing 0, 100, 200, 500, 1,000 and 2,000 pmol free phosphate and 1X reaction buffer in a 50 μl reaction volume were used to generate a standard curve. The activities of protein serine/threonine phosphatases PP-2A, PP-2B, and PP-2C in the extracted proteins were measured using a Synergy H4 Plate Reader. The enzyme activities were quantified based on the standard curve, following the manufacturer’s protocol.

### Multiplexed IHC and quantification

Multiplexed IHC (mIHC) staining was performed on melanoma FFPE tissue samples. Briefly, the slides were first deparaffinized in xylene and then subjected to microwave treatment for epitope recovery. Hematoxylin and eosin (H&E) staining was performed for histopathological evaluation. Subsequently, multiplexed IHC staining was carried out using the Opal 7-Color IHC Kit (PerkinElmer, Cat# NEL811001KT). For melanoma patient specimens, the antibody panel included anti-KDM5B (Sigma-Aldrich, Cat# HPA027179), anti-MITF (Sigma-Aldrich, Cat# HPA003259), anti-SOX10 (Sigma-Aldrich, Cat# HPA068898), and anti-NFKBIE (Sigma-Aldrich, Cat# HPA005941). For melanoma mouse xenografts, the antibodies used were anti-KDM5B (Novus, Cat# NB100-97821), anti-S100B (Dako, Cat# GA504), anti-SOX10 (Cell Marque, Cat# 383A-74), and anti-N-Cadherin (Abcam, Cat# ab18203). The staining protocol followed the PerkinElmer Opal staining kit manual and previous studies^63^. Finally, DAPI (PerkinElmer) was used to stain cell nuclei. Images were acquired using a Vectra Polaris Multispectral Imaging System (PerkinElmer) for whole-slide scanning. inForm Image Analysis software (inForm 2.4, Akoya Biosciences) was used to process and analyze all images. ImageJ was employed to quantify the fluorescence intensities of single cells within the mIHC images.

### ROS production assay

M397 cells were treated with DMSO or 3 μM vemurafenib, HCC827 cells were treated with DMSO or 2 μM erlotinib, and HT-29 cells were treated with DMSO or a combination of 50 μg/mL cetuximab and 1 μM dabrafenib for 3 days. After treatment, cells were collected to assess ROS production. ROS levels were quantified using the ROS-Glo™ H_2_O_2_ assay kit (Promega, Cat# G8820) following the manufacturer’s protocol. Luminescence signals were measured with a Synergy H4 Plate Reader.

### RNA-seq analysis

Reads were aligned against the human genome (hg19) using either TopHat2^64^ or STAR (v2.7.3a)^65^. Read counts were quantified using htseq-count^66^, utilizing known gene annotations from UCSC^67^ with anti-sense (AS) genes removed. Fold-change values were calculated from Fragments Per Kilobase per Million reads (FPKM) normalized expression values, which were also used for visualization following a log2 transformation. Aligned reads were counted using GenomicRanges^68^. Separate comparison p-values were calculated from raw counts using limma-voom^69^, and false discovery rate (FDR) values were calculated using the method of Benjamini and Hochberg^70^. Prior to p-value calculation, genes were filtered to include only transcripts with an FPKM expression level of 0.1 (after a rounded log2-transformation) in at least 50% of samples^71^. Genes were defined as differentially expressed if they had a |fold-change| > 1.5 and FDR < 0.05. To inspect the similarity of the transcriptome of M397 in different time points, we applied consensus clustering using the R package of ConsensusClusterPlus^72^ to define clusters. The top 3,000 most varying genes were used for consensus clustering with the hierarchical clustering method.

### Information-theoretic surprisal analysis

Surprisal analysis was initially formulated to understand the dynamics of nonequilibrium system and has been extended to characterize and simplify complex biological processes^24,26,28^. The proposition of this theoretical approach is that the ensemble of biomolecules in a biological system is subject to the same physicochemical laws as inanimate nonequilibrium systems in physics and chemistry. It approximates quantum state distributions of molecular species within a cell’s molecular ensemble to apply the procedure of maximum entropy to describe a cellular system^24^. Briefly, for a system characterized by a kinetic series of transcriptome measurements, the entropy of the mixture of transcripts in the cell in information theoretic units per a total of *X* transcript is given by

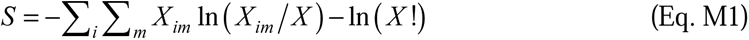

where *X_im_*is the number of transcript *i* that are in the quantum state *m*. *X* = ∑*_i_* ∑*_m_ X_im_* . By defining *x_im_* as the fraction of all the transcript *i* that are in the quantum state *m*, and using Stirling’s approximation, the entropy per *X* transcripts in Eq. M1 can be written as

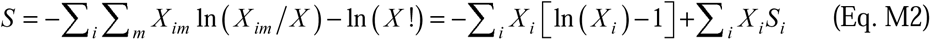

where *X_i_* = ∑*_m_ X_im_* denotes the total number of transcript *i* and *S_i_* = −∑*_m_ x_im_* ln ( *x_im_*) denotes the entropy of a transcript *i*. Eq. M2 describes two contribution terms of the entropy, namely the entropy of mixing and the weighted sum of the entropy of the distribution of quantum states for each of different transcript species.

Resolving the distribution of quantum states requires the partition function for the biomolecules. While this function is not available at the present state of knowledge, for biomolecules with a wide amplitude of structural fluctuations and a large number of low energy conformers, a permissible approximation can be made^73^. Here, we simplify it by assuming two types of quantum states with those of low energy equally populated and those of higher energy not occupied. Then *x_im_* = 1*Q_i_* for state *m* of low energy and *x_im_* = 0 for state *m* of high energy. With this approximation, the entropy in Eq. M2 can be written as

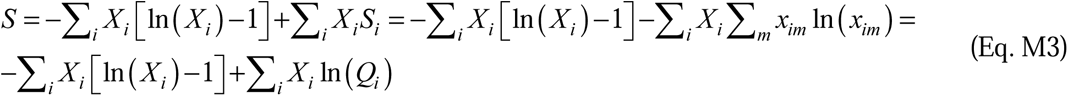

Then, the variation of the entropy can be given by

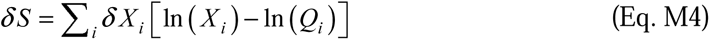

To apply the maximum entropy formalism as a practical tool for a cell at room temperature, we note that, at any point in time, the non-equilibrium cellular system is in a state of maximal entropy subject to constraints. Constraints keep the entropy from reaching its global maximal value by limiting the range of possible variations of the transcript molecule number *X_i_*. This means that the quantity 〈*G_j_*〉 is constant where 〈*G_j_*〉 = ∑*_i_ G_ij_ X_i_* . Here *j* is the label of the constraint and *G_ij_* is the value of the constraint *j* for the transcript *i*.

To seek the maximal entropy state subject to constraints, we use the method of Lagrange multipliers. The Lagrangian of the non-equilibrium system can be written as

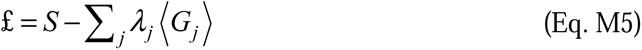

where λ*_j_* is the Lagrange’s multiplier for the constraint 〈*G_j_*〉 = constant. The change of the Lagrangian δ£ due to an unconstrained variation δ*X_i_* has to vanish, which means*δ* £ = *δ S* − ∑ *_j_ λ _j_G_ij_* ∑*_i_ δ X_i_* = 0 . Applying Eq. M4 to this equation, we can obtain

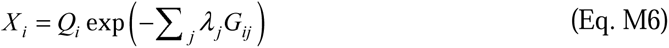

At the unconstrained global stable state, the system reaches the global maximum of the entropy so that the expression level of each transcript is no longer changing over the time scale of interest, namely δ*X_i_* = 0 for all values of *i*. Then we have the Lagrangian £ = *S* − ∑*_i_ γ _i_ X_i_* and γ*_i_* is the Lagrange’s multiplier associated with the constraint *X_i_* = constant. Following the same method above, the distribution of transcripts at the global maximum of entropy can be given by *X_i_^GM^* = exp (*γ_i_*)*Q_i_*, indicating that the population of each gene is proportional to its partition function as the unconstrained global maximum of entropy. Therefore, we can rewrite Eq. M6 as

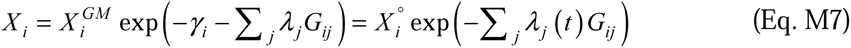

where *X_i_°* denotes the time-independent global stable state and can be determined by the effective number of quantum states for transcript *i*. The exponential term describes the deviation from the global extremum due to the existence of constraints, while the multiplier λ*_j_(t)* is only the time-dependent part of the exponential term. The logarithm of Eq. M7 gives rise to Eq. 1 in the Result section. Surprisal analysis simplifies the transcriptome dynamics into a set of major constraints (or gene modules in biological terms) in addition to the time-invariant global stable state. The global stable state reflects the biological processes that are conserved despite conditions and time points. Each deviation term is a product of a time-dependent multiplier λ*_j_(t)* (the amplitude of a constraint) describing the extent of deviation from the maximal entropy due to the particular gene expression pattern *j* at time *t*, and constraint *G_ij_* describing the time-independent extent of participation of transcript *i* in the gene expression pattern *j*. So, we denote λ*_j_(t)* as gene module scores and *G_ij_* as module-specific contribution score of gene *i* for biological understanding. Gene *i* that displays large positive or negative contribution to a module *j* (high positive or negative *G_ij_*value) represents a gene that is functionally positively or negatively correlated with the module *j*. In other words, the biological function of the module *j* could be inferred by the enrichment analysis of genes with positive and negative G*_ij_* values.

In practice, we determine the global stable state *X_i_°* , the Lagrange multipliers λ*_j_(t),* and constraints by fitting Eq. M7 (or Eq. 1) to the time-series transcriptome data. Singular value decomposition (SVD) has been used to implement this numerical analysis^74^. Briefly, the transcriptome matrix of *k* transcripts measured across *n* time points can be written as **Y** whose entries are the logarithm of the transcript levels, *Y_i_* (*t*) = ln _(_ *X_i_* (*t*)_)_, *i* = 1, 2, …*k*, *t* = 1, 2, …*n* (*k* >> *n* in general). SVD can decompose this non-square matrix by solving the following three matrix equations, namely **Y***^T^* **YP***_j_* = *ω_j_*^2^**P***_j_*, **YY***^T^* **G***_j_* = *ω*^2^**G***_j_*, and **Y= G**Ω**P***^T^* , where **P***_j_* is the eigenvector of *n*-1 components, **G***_j_* is a column vector of *k* components, Ω is a diagonal *n* by *n* matrix whose elements along the diagonal are the eigenvalues ω*_j_*, and *λ _j_* (*t*) = *ω _j_ P _j_* (*t*) . By ranking the eigenvalues in descending order, we can approximate the time-series transcriptome data as a sum with major constraints (or gene modules) as ln 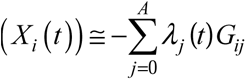, where −*λ*_0_*G_i_*_0_ = ln (*X_i_*^°^) and *A* is the smallest number of constraints that can provide a good approximation for the transcriptome data.

Applying the surprisal analysis to our time-series transcriptome data simplifies the transcriptome dynamics into two major gene modules in addition to the time-invariant global stable state. The sum of the global stable state and the first two gene modules recapitulates the time-series transcriptome data (Fig. 2a). The time-dependent module scores of the first two modules depict the global transcriptome trajectories of cells upon drug treatment and removal, as shown in the cyclic trajectories of Fig. 2b. To project the transcriptomes of other melanoma cell lines onto the two-module defined space, the module-1 and module-2 scores of other published datasets on melanoma cell lines^5^ were calculated using the same approach (Fig. 5b). Cell lines projected onto the similar region of long-term BRAFi-treated M397 cells will be transcriptionally more similar to the BRAFi-resistant transcriptional state of M397 cells.

### Visualization of gene modules using self-organizing mosaic maps

To visualize the natural log-transformed transcriptome dataset and contributions from each gene module λ*_j_(t)G_ij_*calculated from surprisal analysis, self-organizing mosaic maps (SOMs) were used. The SOMs provided a high-resolution visualization of patterns by plotting individual samples as single 2-dimensional heatmaps. Thousands of input genes were assigned to 625 rectangular “tiles” (SOM nodes), each representing a mini-cluster of genes. These mini-clusters were arranged to form a pattern within a 2-dimensional mosaic map on the SOM grid. Each mini-cluster of genes was mapped onto the same tiles in every map, with the color of each tile representing the relative average expression of the gene mini-cluster within that tile. Thus, tiles at the same location represented the same group of genes across different conditions. Clusters with the most similar kinetics over the transition were placed adjacent to each other in the mosaic map, enhancing the visualization of gene expression dynamics. The Gene Expression Dynamics Inspector (GEDI) package was utilized to implement the SOM visualization^30^.

### Dynamic system modeling of the gene module interactions

The time-resolved transcriptome data can be deconvoluted into a global stable state and two time-varying gene expression modules, each summarized by a weighted sum (“score”, or [gene_Mj_ ]) of constituent gene expressions:

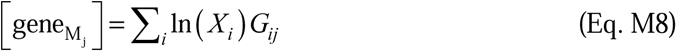

where *X_i_* is the expression level of gene *i* included in the module *j*, and *G_ij_* is the relative contribution of the gene *i* to the gene module *j*. To quantitatively understand the interactions of the two gene modules involved in the reversible cell state dedifferentiation, we determined a mathematical relationship between the averaged expression levels of module-associated genes [gene_Mj_ ] through a set of coarse-grained ordinary differential equations (ODEs) based on the first-order mass equations shown to be physiologically plausible and accurately descriptive of cancer cell state transition at the cell population-level^75^. Such a system may lead to the maintenance of multiple stable cellular states, and the relative stability of these states may be perturbed by environmental perturbations^75,76^. Our experimental observations are consistent with such a model.

If we consider a set of *n* gene module scores, we can describe them as a vector **g** with *n* dimensions. We assume that the component gene module scores are the principal regulators of themselves and of each other; time-dependent changes in the gene module scores due to module-module interactions can be summarized by an *n* × *n* matrix **M**. We also assume that gene modules are additionally affected by basal levels of transcriptional synthesis or degradation with respect to time, denoted by an *n*-dimensional vector **b**. Then, the time-dependent change in the gene module scores may be expressed as

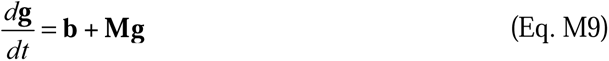

Applying this equation to our specific context, let 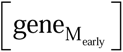 and 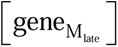 denote the averaged expression levels of the top 500 genes that have the highest absolute positive or negative *G_ij_* values associated with M_early_ and M_late_, respectively. The changes in 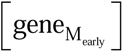 and 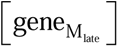 with respect to time are determined by constitutive synthesis or degradation (*B_e_* and *B_l_*), first-order auto-regulation (*M_e-e_* for M_early_ and *M_l-l_*for M_late_), and first-order regulation imposed by the other gene module (*M_e-l_* for M_early_ regulating M_late_; *M_l-e_* for M_late_ regulating M_early_). The resulting ODE system of Eq. M9 can thus be expanded as

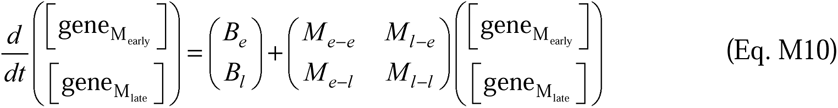

More specifically, for drug treatment condition, we have *G_i1_*-positive genes and *G_i1_*-negative genes (genes that are positive or negatively correlated with M_late_), which are paired with *G_i2_*-positive and *G_i2_*-negative genes (genes that are positive or negatively correlated with M_early_) respectively. Therefore, we have 4 different scenarios for the drug treatment condition. Similarly, we also have 4 different scenarios for the drug removal condition. For Eq. M10 (essentially Eq. 2 in the Result section), we fitted all coefficients simultaneously using a Markov Chain Monte Carlo (MCMC) simulation. To simulate the resulting trajectories following Eq. M10, we applied the forward Euler method, incrementing the score of each gene module with respect to time as

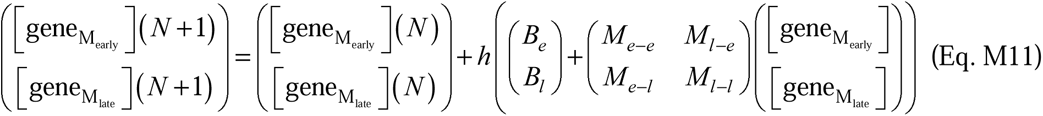

where *N* is the step number (i.e., time increment) starting from day-0 and *h* is the step size, which we set to 0.5 days for fitting.

We assumed Gaussian distribution of coefficient probabilities. All coefficients were initialized using independent, random uniform distributions, assuming each coefficient had uniform prior probabilities. Thus, the log-posterior probability used in the MCMC simulation was

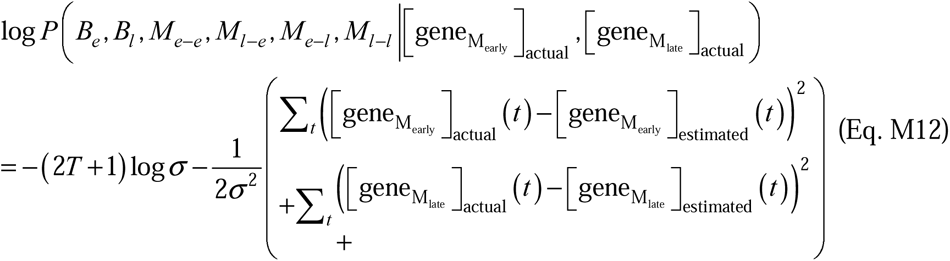

where *T* is the total number of calculated time points and σ is the estimated variance; σ was initialized using an exponential distribution assuming a Jeffreys prior.

We also constrained the simulation such that fits leading to physiologically implausible oscillations of 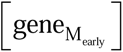 or 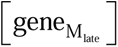, namely frequencies greater than the Nyquist frequency (1 / day) of our experimental sampling, were considered invalid fits and discarded. Assuming that the last experimental observation of expression represented the steady state, we extended the fitting data by up to three hypothetical observations that had values identical to the last point of the experimental data (Extended Data Fig. 3b). These hypothetical data points were each spaced four days apart after the last true experimental point, thereby allowing the fits to recognize this experimental steady state.

We fitted coefficients for the above system for both the forward (dedifferentiation) and reverse (re-differentiation) transition trajectories (Extended Data Fig. 3 and Extended Data Table 1), the latter of which was measured by removing drug from the cell culture after 29 days of drug treatment.

### Gene set enrichment analysis

Gene Set Enrichment Analysis (GSEA) was performed using GSEA v2.2.3 software with 1,000 permutations and weighted enrichment statistics. The normalized enrichment score (NES) was calculated for relevant gene sets curated in the Molecular Signatures Database (MSigDB). The MITF signature gene set was constructed based on a previous report^77^, and the KDM5B target genes set was constructed using the CHEA Transcription Factor Targets database^78^. For single-sample gene set enrichment, as shown in Extended Data Fig. 1d, we used the GSVA program to derive the absolute enrichment scores of relevant gene signatures curated in MSigDB C2. The normalized log2 RPKM values were used as input for GSVA in the RNA-seq mode.

### Transcription factor target and motif enrichment analysis

To identify the driving transcription factors in the module-2 (M_early_) process, we employed two different approaches. In the first approach, we filtered transcription factors (TFs) and co-factors associated with module-2 by selecting those with a Pearson correlation coefficient, |r|, greater than 0.8 with the module-2 amplitude (λ_2_). These were defined as module-2 associated TFs/co-factors. We then acquired the downstream target genes for all module-2 associated TFs/co-factors using the public database TFtargets (https://github.com/slowkow/tftargets). The KDM5B gene targets were manually verified using ChIP-seq data from ENCODE database (GSE101045). Next, we further filtered the module-2 associated TFs/co-factors based on the overlap of their downstream target genes with genes associated with module-1 (M_late_) (|r| > 0.8 with module-1 amplitude (λ_1_)). Specifically, a TF in module-2 was selected as a candidate driving TF if its downstream target genes were over-represented in the module-1 process (Hypergeometric test with Bonferroni correction, FDR ≤ 0.05). In the second approach, we used HOMER to identify enriched motifs in the promoter sequences of genes associated with module-1 (|r| > 0.8 with module-1 amplitude (λ_1_)). The parameters used were: -len 6,8,10,12, -start -1800, -end 100, -b, and -mset vertebrates. Potential TFs were then inferred based on the enriched motif information (binomial distribution, p-value < 0.05).

### ChIP-seq analysis

Reads were mapped to the human genome hg19 using bowtie2^79^. Identical aligned reads were deduplicated to avoid PCR duplicates. Peaks were called on the merged set of all ChIP-seq reads from M397 cells using MACS2 with the following parameters: --nomodel, --broad^80^. Peaks were assigned to the nearest gene with the closest transcription start site (TSS). Peaks from each sample were compared and merged. Read counts for each sample were measured using the FeatureCounts tool from Subread v2.0.3^81^. Differential analysis between D0 and any other samples (D3, D32, DR30) was performed using DESeq^82^. Differential binding regions were identified if the absolute log2 fold change was greater than 1 and FDR < 0.05. To visualize peaks in each sample, bedGraph files were generated using MACS2 with following parametes: --nomodel, --broad, --bdg, --SPMR. The bedGraph files were then converted into bigWig files using the bedGraphToBigWig tool. ChIP-seq signals within the promoter regions of RelA target genes or genome-wide were visualized using deepTools v3.0.2^83^. To assess changes in baseline histone mark profiles across multiple melanoma cell lines, the average histone ChIP-seq signals at the promoter regions of RelA target genes were calculated by averaging the normalized total read counts of H3K4me3 and H3K27ac marks within the TSS ± 2 kb regions across the 307 RelA target genes.

### ATAC-seq analysis

Adaptor sequences were first trimmed from the reads using Cutadapt. The reads were then aligned to the hg19 genome using Bowtie2 with standard parameters and a maximum fragment length of 2,000^79^. Identical aligned reads were deduplicated to avoid PCR duplicates, and the deduplicated reads were filtered for high quality (MAPQ ≥ 30). Peaks were called on the merged set of all ATAC-seq reads from M397 using MACS2 with following parameters: --nomodel, -broad, -q 1e-5^80^ and filtered to remove putative copy number varied regions^84^. Differentially accessible regions between Day 0 (D0) and other samples (D3, D32, DR) were identified using diffReps with a window size of 500^85^. Differential binding regions were defined if the absolute log2 fold change was greater than 1 and FDR < 0.05. These regions were then compared and merged with ATAC-seq peaks called by MACS2. To visualize peaks in each sample, the same routine in ChIP-seq analysis was applied. ATAC-seq profiles of differentially accessible region M397 samples were generated by using ngs.plot.r with following parameters: -G hg19 -R bed -L 1000 -GO km -KNC 4 -SC 0,3.5. The profiles of unchanged ATAC-seq peaks in M397 samples were plotted by using ngs.plot.r with following parameters; -G hg19 -R bed -L 1000 -GO total - SC 0,3.5. HOMER was used to identify over-represented motifs in the set of differentially accessible peaks, using a background set of peaks that did not significantly change, and using the parameters: “-size given -len 6,8,10,12 -mset vertebrates -bg”^84^.

ATAC-seq signals within the promoter regions of RelA target genes were visualized using deepTools v3.0.2^83^. Read counts were calculated around the differential peaks (± 1 kb) with a window size of 10 bp and normalized by RPKM. RPKM (per bin) = number of reads per bin / (number of mapped reads (in millions) * bin length (kb)). To assess changes in baseline chromatin accessibility profiles across multiple melanoma cell lines, the average ATAC-seq signals of RelA target genes for each cell line were calculated by averaging the normalized total read counts of all peaks within ± 2 kb flanking the TSS to TES regions across the 307 RelA target genes.

### Integrated analysis of time-series multi-omics data

To identify RelA target genes modulated by RelA-mediated chromatin remodeling and transcriptional repression, we developed an analytical pipeline integrating time-series transcriptome, ATAC-seq, and ChIP-seq data across the dedifferentiation transition. First, ChIP-seq and ATAC-seq peaks were called using SICER2 with standard parameters^86^. Differential peaks between D0 and other time points (D3, D32) were assessed using sicer_df with default parameters and FDR<0.01. Target genes were identified whose promoter regions (TSS ± 2 kb) exhibited increased RelA binding peaks and overlapping reductions in H3K4me3 and H3K27ac peaks at D3 using bedtools v2.27.1. These genes were further filtered based on a consequent reduction in chromatin accessibility at the promoter regions and decreased expression levels with fold changes greater than 2 at D32. This analysis identified 307 RelA target genes involved in RelA-mediated transcriptional repression through the reduction of activation histone marks and consequent reduction of chromatin accessibility upon continued BRAF inhibition. Enrichment analyses of the 307 RelA target genes against curated gene sets were performed using Enrichr^87^. Significantly enriched terms (p < 0.05) were ranked by odds ratios.

To infer downstream transcription factors (TFs) modulated by RelA, we assessed changes in RelA binding profiles, chromatin accessibility, and H3K4me3 and H3K27ac activation histone marks at the RelA binding sites within the promoter regions (TSS ± 2 kb) of all the TFs among the 307 RelA target genes identified. We evaluated the statistical significance of these changes (i.e., differential peaks identified by sicer_df) between D32 and D0 and found that SOX10 displayed the most significant changes for all three epigenetic alterations (Fig. 3h).

### Patient data analysis

Paired patient data before and after BRAFi (or MAPKi) treatments were analyzed to evaluate dedifferentiation-related gene expression levels. The log fold changes in gene expression levels (post-BRAFi vs. pre-BRAFi) were visualized as a heatmap (Extended Data Fig. 2d). These data were sourced from two published papers^16,22^. The original gene expression levels and associated patient identification numbers are provided in Supplementary Dataset 2.

### Statistical analysis

Statistical analyses were conducted using GraphPad PRISM 10 (GraphPad Software, Inc.) unless otherwise specified. Statistical significance between two groups was assessed using a two-tailed Welch’s t-test with a significance threshold of p < 0.05, with multiple comparisons corrected using the two-stage step-up (Benjamini, Krieger, and Yekutieli) method, unless otherwise noted.

## Acknowledgements

We thank Sui Huang (ISB), Lee Hood (ISB), and the Michael Elowitz lab (Caltech) for comments on the manuscript. We acknowledge the following funding agencies and foundations for support: NIH/NCI U01CA217655 (to R.L., A.R., J.R.H., and W.W.), U54CA274509 (to R.L., J.R.H., and W.W.), Andy Hill CARE Fund (to W.W.), Phelps Family Foundation (to W.W.), NIH/NCI P01CA168585 (to A.R.), R35CA197633 (to A.R.); Dr. Robert Vigen Memorial Fund, the Ressler Family Fund, and Ken and Donna Schultz (A.R.); the Jean Perkins Foundation (J.R.H.); ISB Innovator Award (Y.S.). National Key Research and Development Program Grant 2016YFC0900200 (to Q.S.), National Natural Science Foundation of China Grant 21775103 (to Q.S.), 81672696 (to J.G.), and 81772912 (to Y.K.). L.R. was supported by the V Foundation-Gil Nickel Family Endowed Fellowship and a scholarship from SEOM.

## Author contributions

J.R.H. and W.W. conceived the project and supervised the study. Y.S., C.L., X.L., G.L., S.S., R.H.N., S.W., L.R., V.L., V.L., J.C., Z.W., Y.T., H.C., A.H.C.N., S.P., M.X., D.J., and J.W. performed the experiments. C.L., X.L., and C.W. performed bioinformatics analysis. Y.S., X.L., R.L., J.R.H., and W.W. established the computational models. Y.S., C.L., X.L., G.L., J.W.L., C.W., V.L., Z.W., G.Q., J.W., I.S., X.W., Q.S., R.L., A.R., D.B., J.G., J.R.H., and W.W. analyzed the data. Y.K., Y.X., and J.G. contributed the clinical specimens and expertise. Y.S., C.L., X.L., J.R.H., and W.W. wrote the original draft. Everyone reviewed the paper.

## Extended Data

**Extended Data Fig. 1.**
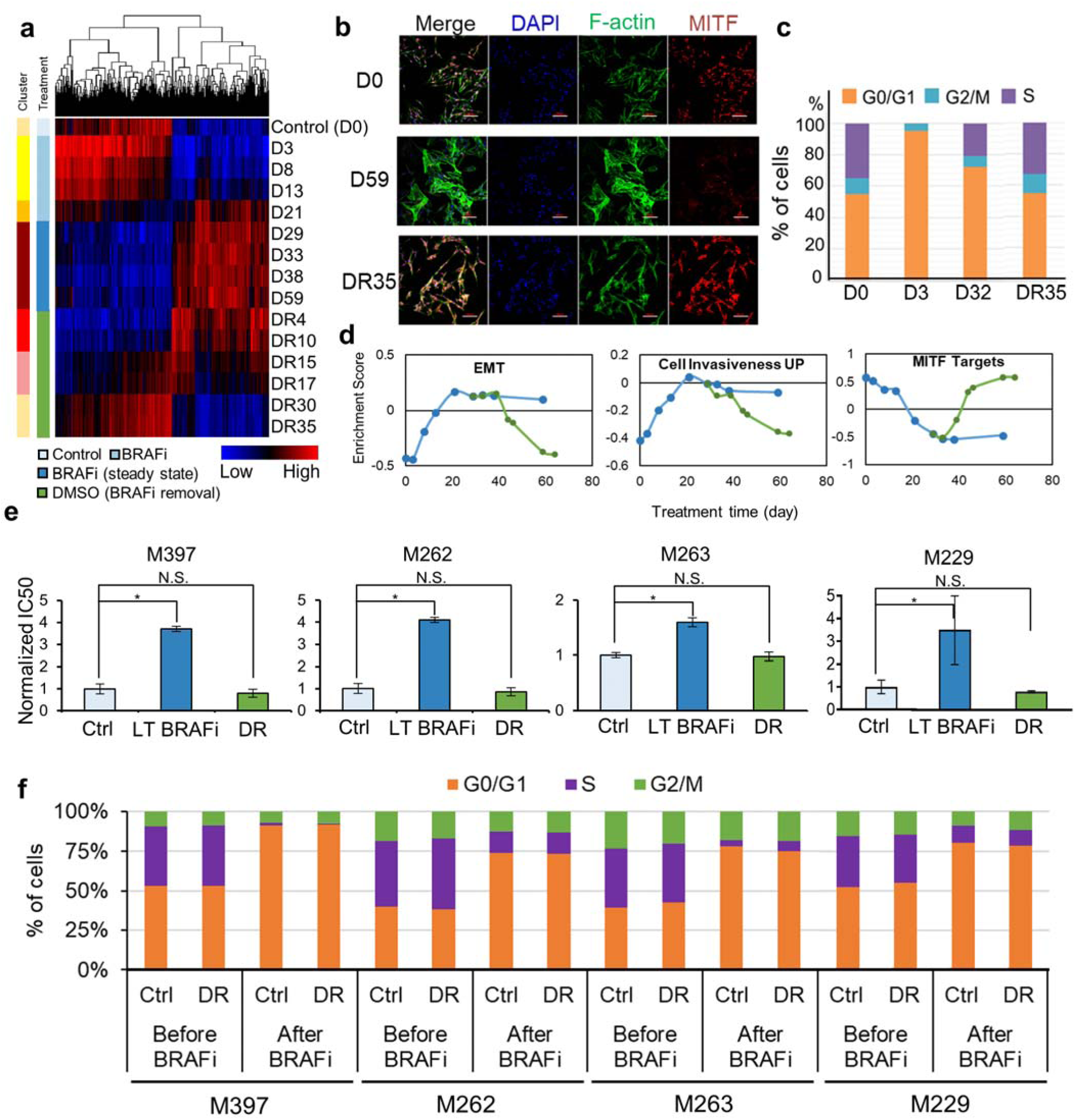
Transcriptome signature, cell cycle, and drug sensitivity associated with the drug-induced dedifferentiation transition and its reversion across *BRAF*-mutant melanoma cell lines. **a.** Heatmap clustering of differential expressed genes (DEGs) over the BRAFi (vemurafenib) treatment and removal for M397 cells. Sidebars denote consensus clustering results of DEGs from the samples (6 clusters) and their treatment conditions. DR30 and DR35 cluster with the control sample (DR: drug removal). **b.** Immunofluorescent staining of M397 cells at different stages of the dedifferentiation transition. M397 cells before treatment (D0, first row), after 59 days of BRAFi treatment (D59, second row), and 35 days after drug removal (DR35, third row) were stained for MITF (red), F-actin (green), and DAPI (blue). The cell morphology at D0 is similar to that at DR35, while the morphology at D59 is distinct from the other two conditions (scale bar 100 μm). **c.** Cell cycle distribution of M397 cells at specified time points across the dedifferentiation transition. Clear cell regrowth with more S phase cells was observed at D32 of BRAFi treatment, confirming the development of adaptive resistance. **d.** GSVA enrichment scores of representative gene sets over the course of the dedifferentiation transition and its reversion. Enrichment scores at different time points are shown as dots connected with solid lines. The scores stabilize after prolonged drug treatment (after 29 days of BRAFi, blue line) and return to the day 0 enrichment score after long-term drug removal (green line). **e.** Increased drug tolerance and reversed drug sensitivity across multiple melanoma cell lines with varying baseline sensitivities to BRAF inhibition, evaluated by IC50 values of vemurafenib. LT: long-term (30 days); DR: drug removal. (Mean ± SD, *p<0.05, compared to control, n=3). **f.** Stacked bar plot showing the fraction of cells viable in G0/G1, S, and G2/M phases (y-axis) for different melanoma cell lines measured either before treatment (Ctrl), or after continuous treatment with BRAFi for 30 days followed by drug removal for another 30 days (DR). Cells at both conditions were retreated with either DMSO (i.e. before BRAFi) or vemurafenib (after BRAFi) for another 3 days for cell cycle analysis. Cells that underwent drug treatment and removal exhibit a highly similar cell cycle distribution as those never received drug treatment.

**Extended Data Fig. 2.**
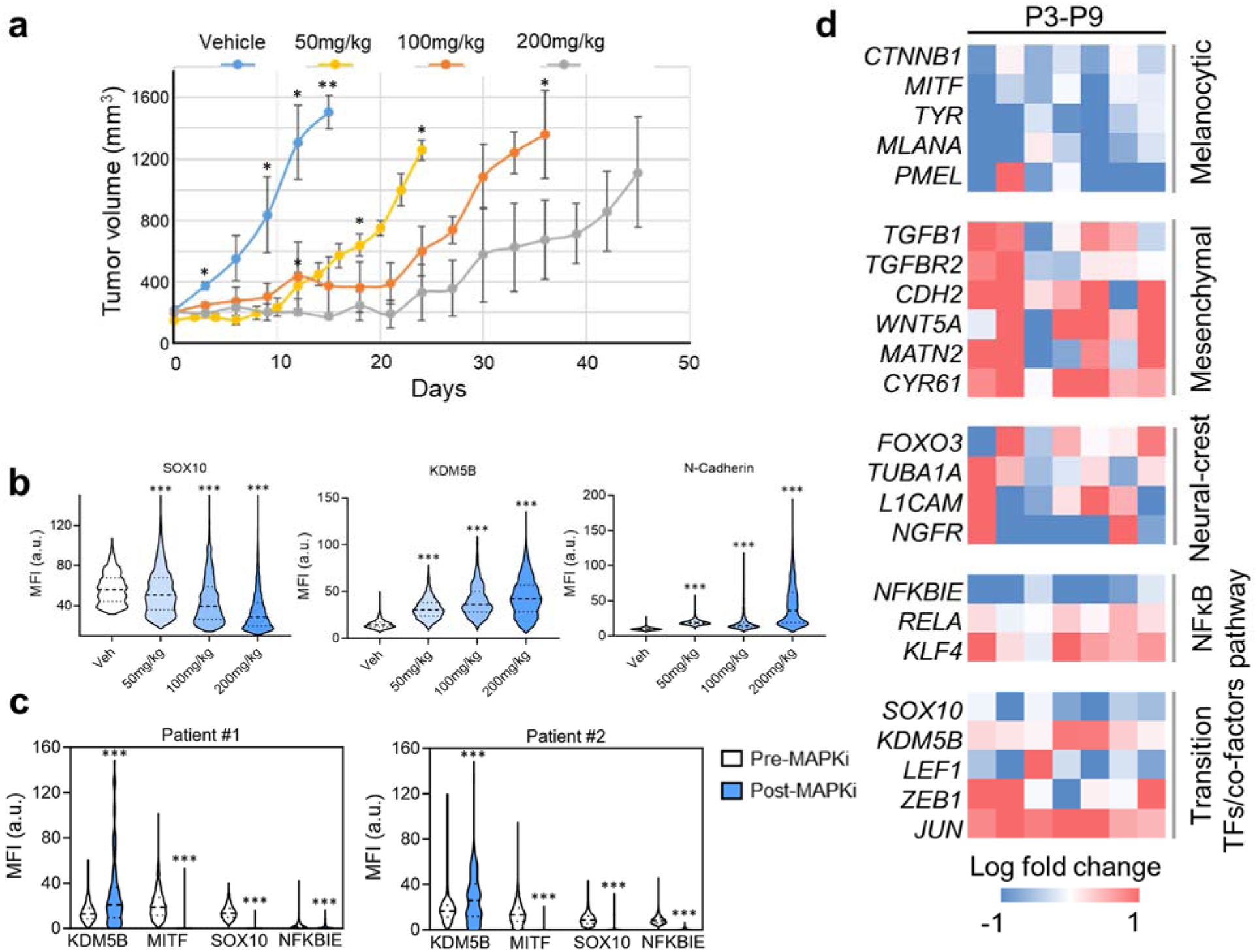
Drug-induced dedifferentiation transition in melanoma mouse xenograft models and patient samples. **a.** Tumor growth curve of an orthotopic mouse xenograft model bearing *BRAF*-mutant melanoma cells treated with increasing doses of BRAF inhibitor vemurafenib (n=3, *p<0.05, **p<0.005 with respect to 200mg/kg group). **b.** Quantification of mIHC staining of the *BRAF*-mutant melanoma mouse xenograft model treated with varying doses of BRAFi, as shown in Fig. 1e. Veh: vehicle. Data are presented as mean fluorescent intensity (MFI) per cell (Kruskal-Wallis test with correction for multiple comparison using Dunn’s test, ***p<0.0005 compared to vehicle control, all the cells in the image are quantified for each treatment condition). **c.** Quantification of mIHC staining for selected markers in two patients before and after BRAFi treatment. Data are represented as fluorescent intensity (MFI) per cell (Two-tailed Mann-Whitney test, ***p<0.0005 compared to respective pre-BRAFi condition, cells within the white dashed lines are quantified for the post-BRAFi condition). **d.** Log-fold change in the expression of dedifferentiation transition-related genes (post-BRAFi vs pre-BRAFi), collated from published datasets of *BRAF*-mutant melanoma patients treated with BRAF inhibitors. See also Supplementary Table 2.

**Extended Data Fig. 3.**
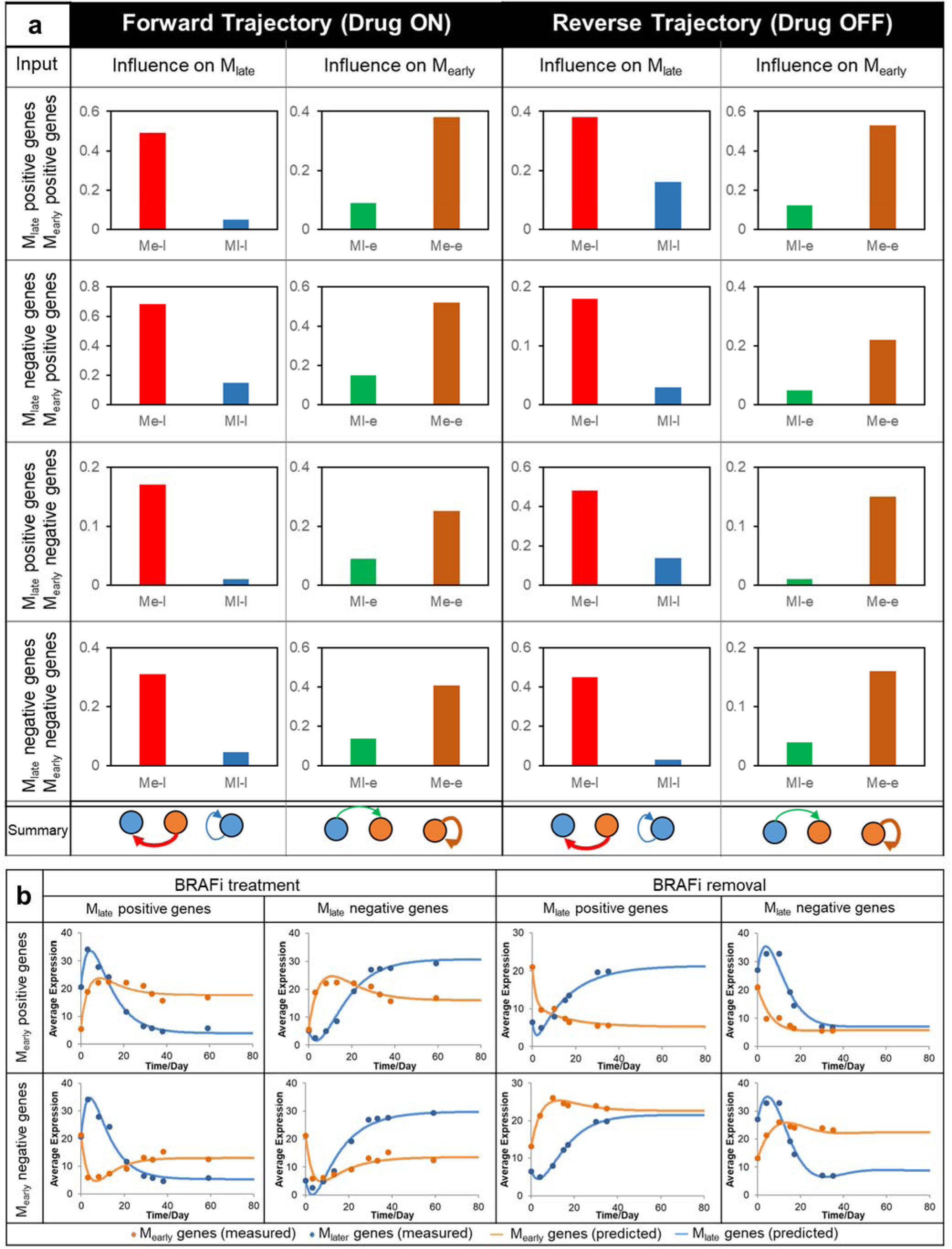
Best fitted parameters and predictions of the dynamic ODE modeling of drug-induced dedifferentiation transition and its reversion in M397. **a.** Module-module interaction coefficients in the ODEs determined by fitting the ODE model to the average expression level of the top 500 genes associated with each gene module. M_late_ positive genes (those positively contributing to M_late_, i.e. *G_i1_*>0) and M_late_ negative genes (those negatively contributing to M_late_, i.e. *G_i1_*<0) are paired with M_early_ positive (those positively contributing to M_early_, *G_i2_*>0) and M_early_ negative genes (those negatively contributing to M_early_, *G_i2_*<0), respectively, forming four different pairs of scenarios for either forward or reverse transition trajectories. **b.** Experimentally measured (dots) and ODE fitted (smooth lines) average expression levels of genes associated with the two gene modules in the forward and reverse directions of the cyclic transition. M_late_ positive/negative genes are paired with M_early_ positive/negative genes, respectively, to form four different pairs of scenarios for either forward or reverse transition trajectories.

**Extended Data Fig. 4.**
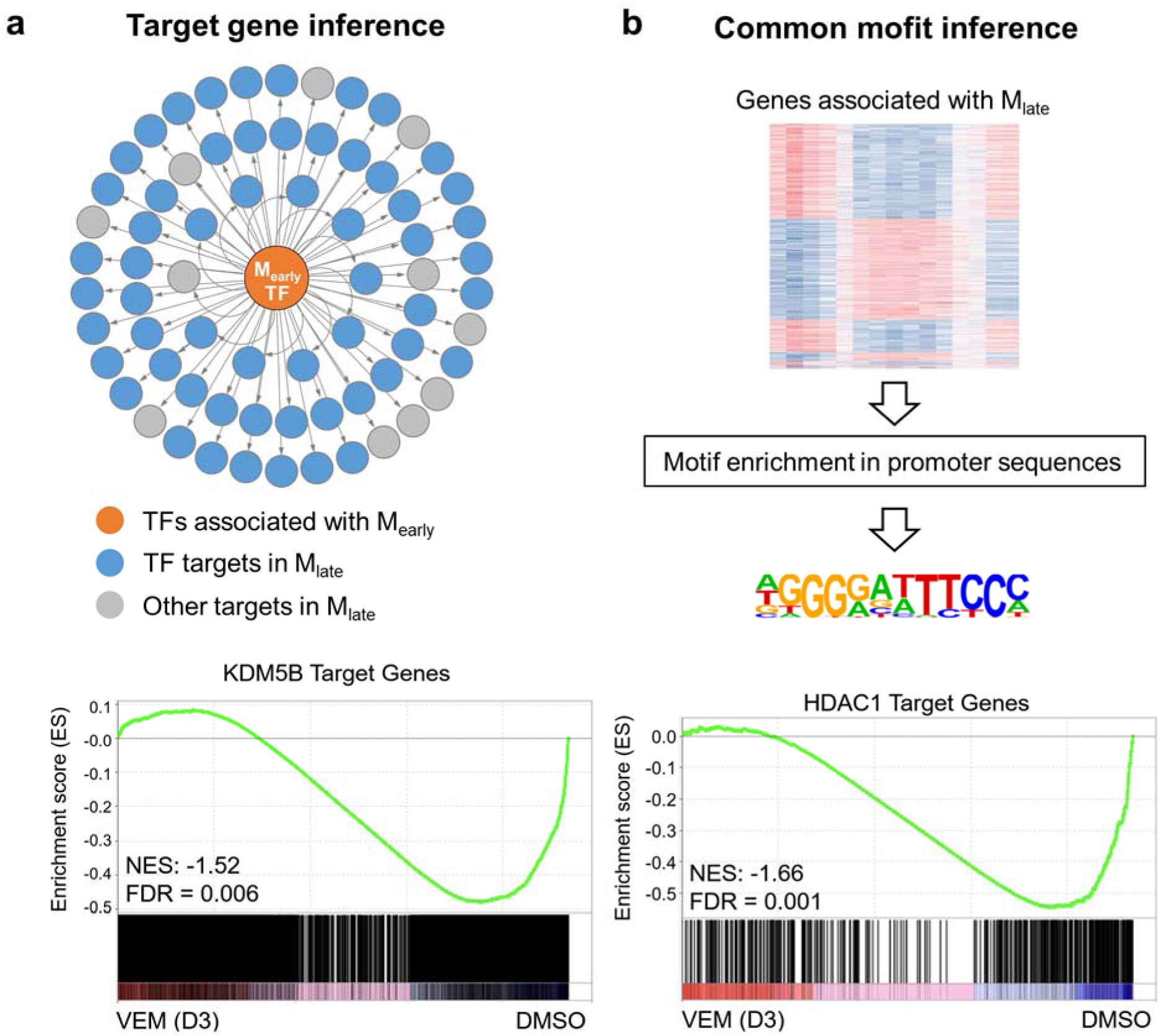
Strategies for inference of critical regulators driving the initiation of the dedifferentiation transition. **a.** Target gene inference based on the dynamic interactions between the two gene modules. The TFs/co-factors whose expression kinetics are highly correlated with the module scores of M_early_ are mapped to their target genes. The enrichment of these target genes among those highly correlated with the module scores of M_late_ is then assessed. Inferred transition-driving TFs/co-factors whose target genes are significantly overrepresented in genes associated with M_late_ are ranked by their absolute correlation coefficients with M_early_ scores. **b.** Common motif inference to extract enriched promoter-binding motifs from genes highly correlated with M_late_. This analysis identifies TFs/co-factors that bind to these motifs and regulate the cell state dedifferentiation. **c.** Gene set enrichment analysis showing significant downregulation (negative enrichment) of KDM5B and HDAC1 target genes upon 3 days (D3) of 3 μM vemurafenib (VEM) treatment compared to the DMSO control. NES: normalized enrichment score.

**Extended Data Fig. 5.**
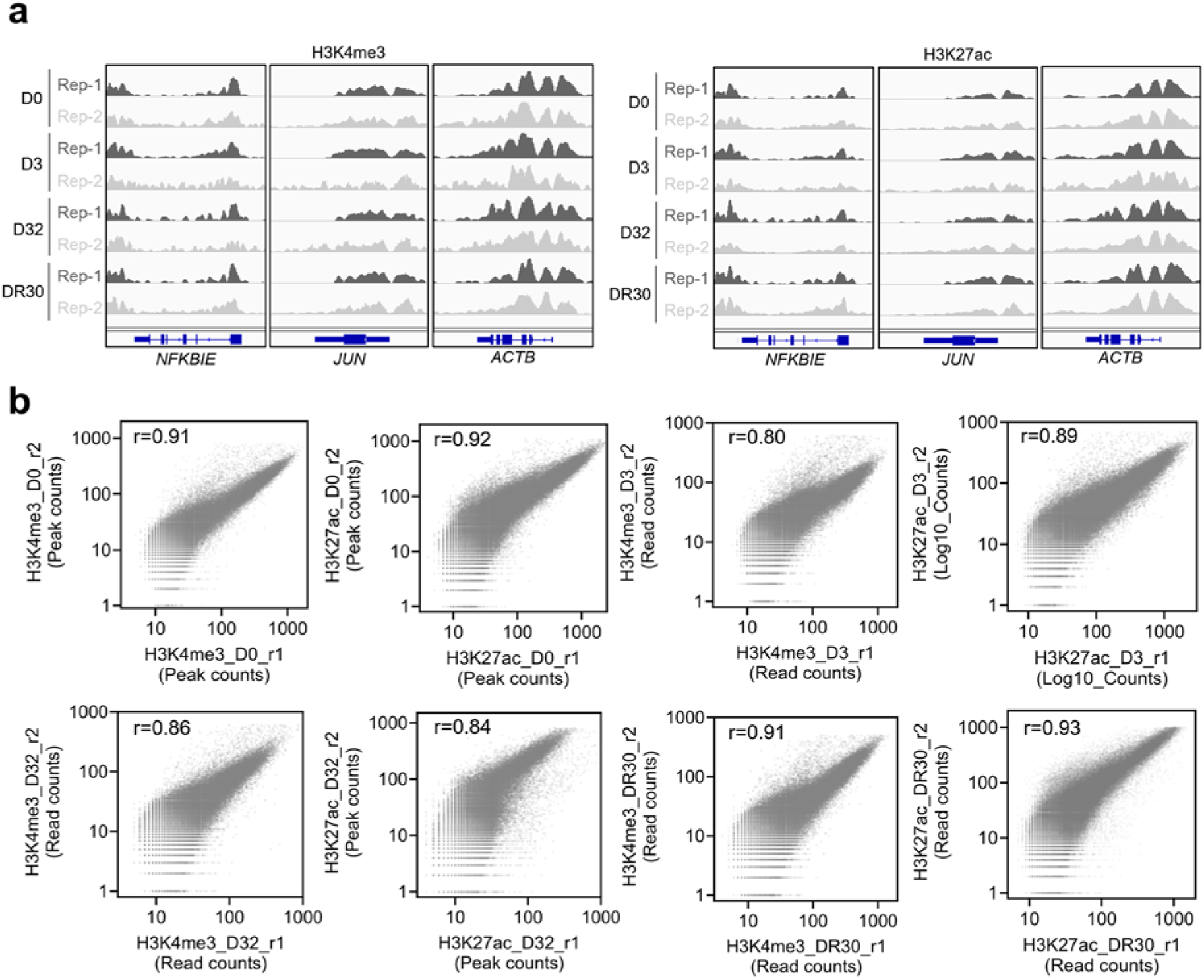
Consistency between ChIP-seq replicates. **a.** Genome browser view of representative histone ChIP-seq profiles of M397 cells at different drug treatment time points during the drug-induced dedifferentiation transition and its reversion, shown across two biological replicates **b.** Scatter plots showing the consistency between two biological ChIP-seq replicates for both H3K4me3 and H3K27ac at different drug treatment time points. Pearson correlation coefficients (r) are listed, with p<0.0001 for all pairwise correlations, indicating high consistency between replicates.

**Extended Data Fig. 6.**
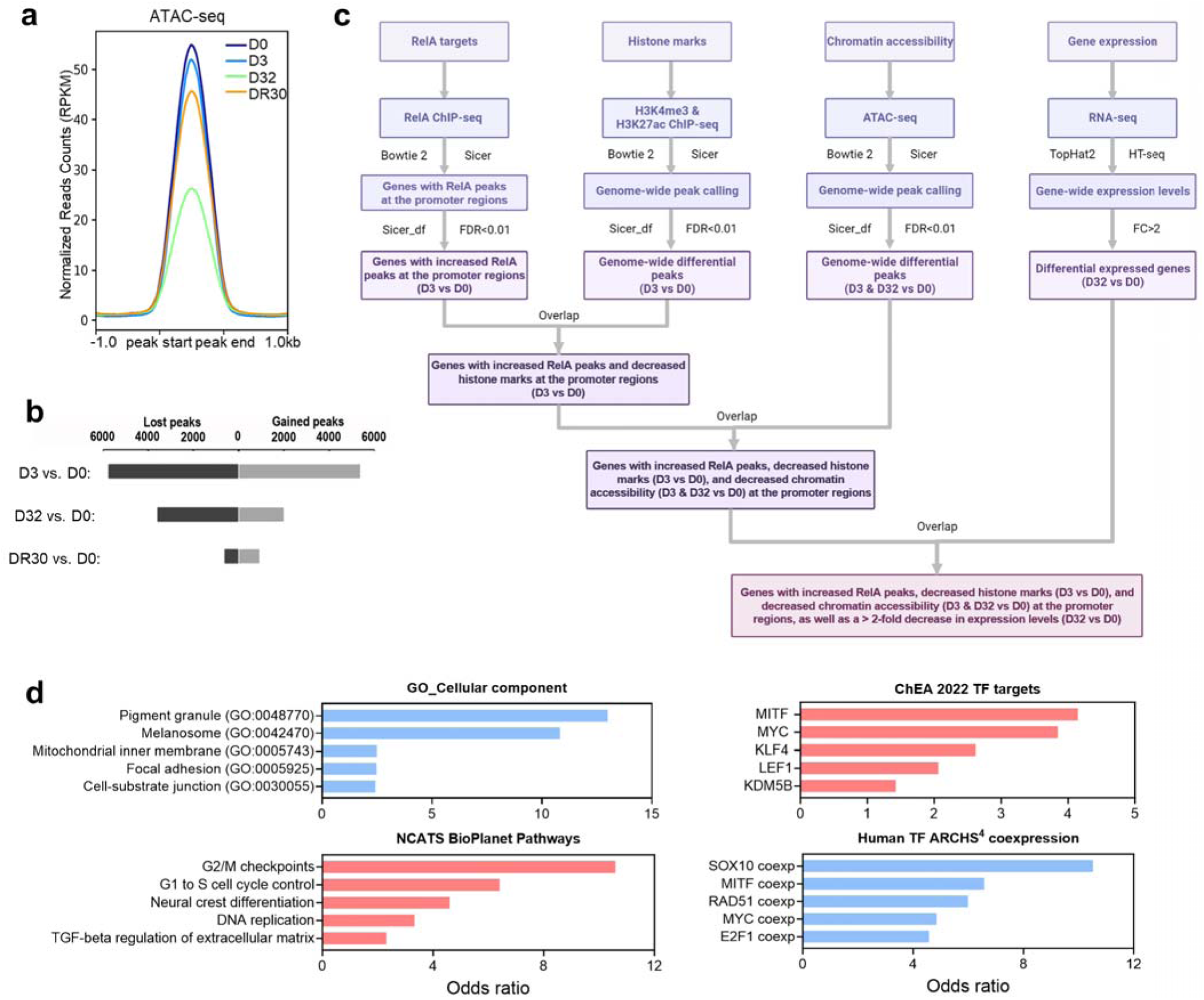
Integrated analysis of time-series multi-omics data during drug-induced dedifferentiation transition. **a.** Chromatin accessibility change assessed by average peak signal of ATAC-seq across the entire genome. The x-axis shows flanking regions of ± 1kb around the peak region. **b.** Differential cis-regulatory element profiles assessed by ATAC-seq at specified time points across the dedifferentiation transition and its reversion. **c.** Illustration of the integrated multi-omics analysis pipeline used to pinpoint RelA target genes whose expression levels are strongly associated with RelA-mediated chromatin remodeling and transcriptional repression. **d.** Representative enriched programs of the 307 RelA target genes within specified molecular signature databases (p < 0.05).

**Extended Data Fig. 7.**
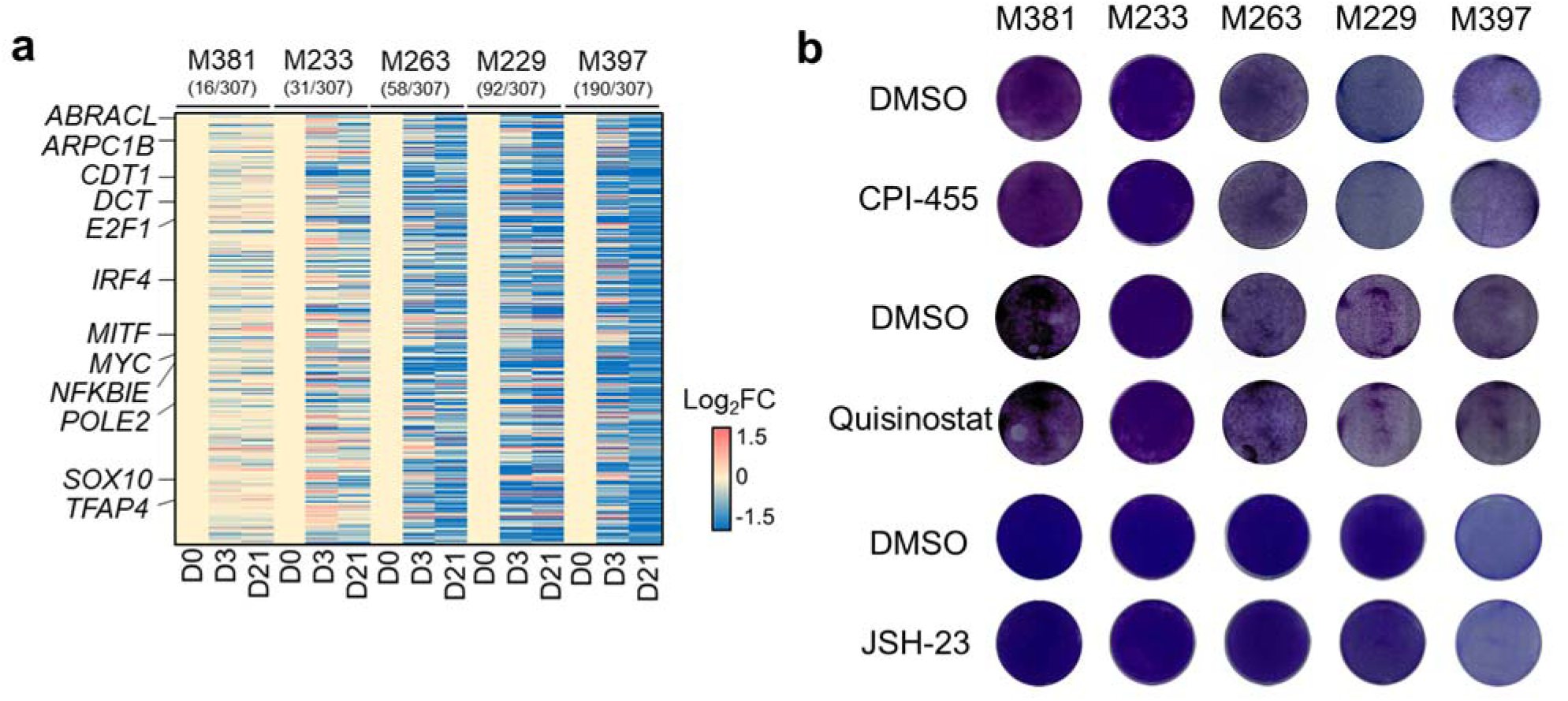
Epigenetic plasticity across different melanoma cell lines. **a.** Heatmap showing the Log2 fold changes (Log_2_FC) in expression levels of the 307 RelA target genes at different time points of VEM treatment relative to untreated control (D0) across different melanoma cell lines. The numbers in the parentheses under each line indicate the number of genes that displayed at least a 2-fold reduction in expression levels at 21 days of drug treatment, a mid-transition time point during the drug-induced dedifferentiation process. **b.** Clonogenic assay of cells treated with KDM5 inhibitor (CPI-455), HDAC inhibitor (Quisinostat), and RelA nuclear translocation inhibitor (JSH-23) or DMSO for the same duration across a panel of melanoma cell lines, showing no significant toxicity to the cells when used alone at the doses used in combination therapies in Fig. 6e, compared to DMSO control.

**Extended Data Fig. 8.**
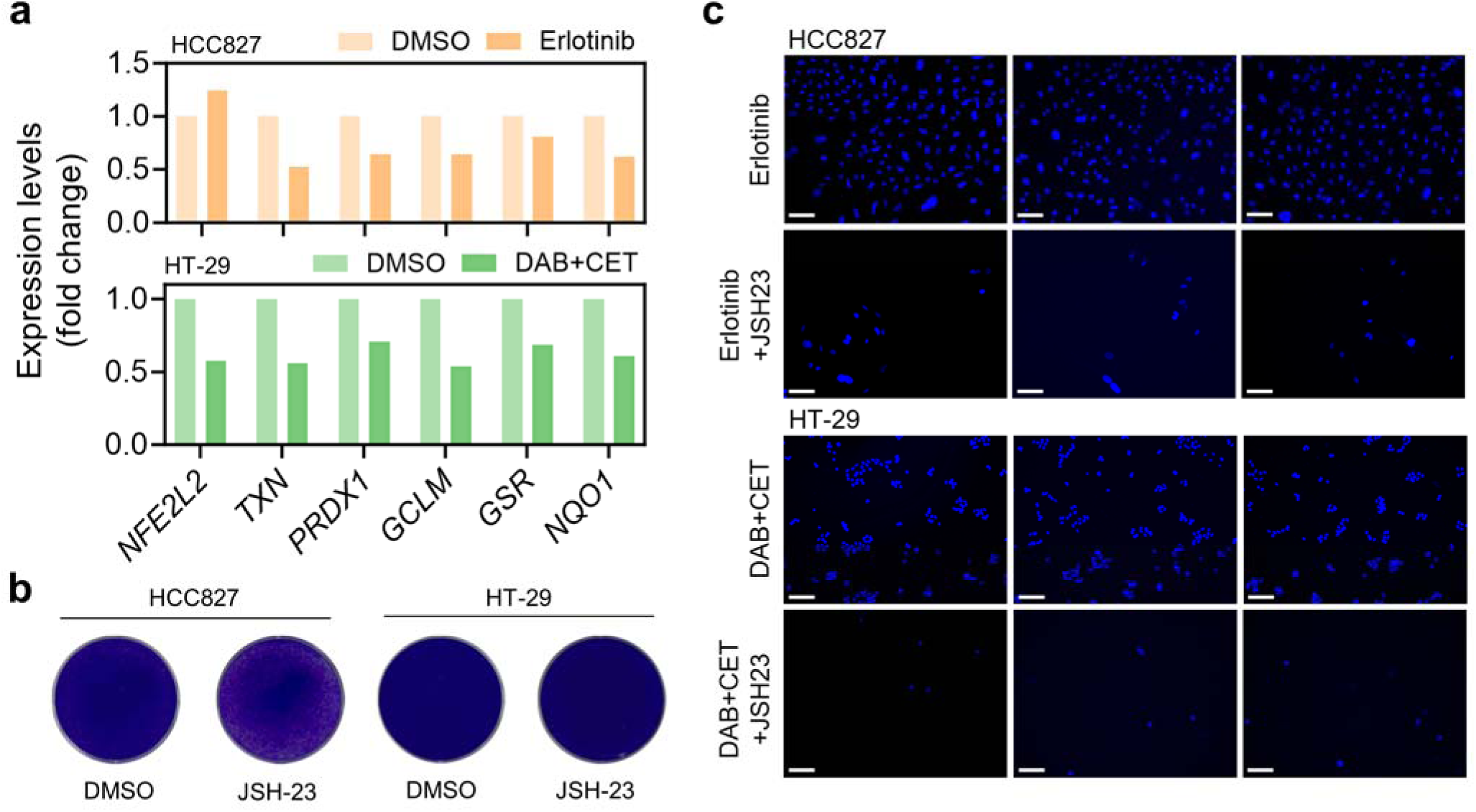
Oxidative stress-mediated NF-κB/RelA activation in the transition towards DTP states in other cancer types. **a.** Changes in expression levels of critical genes associated with antioxidant defense after 3 days of erlotinib (2 μM) treatment for HCC827 cells or DAB (1 μM) + CET (50 μg/mL) for HT-29 cells compared to DMSO controls. **b.** Toxicity of JSH-23 was assessed by comparing cells treated with JSH-23 or DMSO control for the same duration, showing no significant toxicity of JSH-23 to the cells when used alone at the doses used in combination therapies in Fig. 7m and 7n, compared to DMSO control. **c.** Representative fields of view showing Hoechst nuclear staining images. These images illustrate the relative viability of HCC827 cells treated with erlotinib alone or in combination with RelA nuclear translocation inhibitor (JSH-23), as well as HT-29 cells treated with DAB+ CET alone or in combination with JSH-23. Treatment was stopped upon clear cell regrowth in the erlotinib or DAB+CET group (scale bar: 50 µm)

**Extended Data Table 1.**
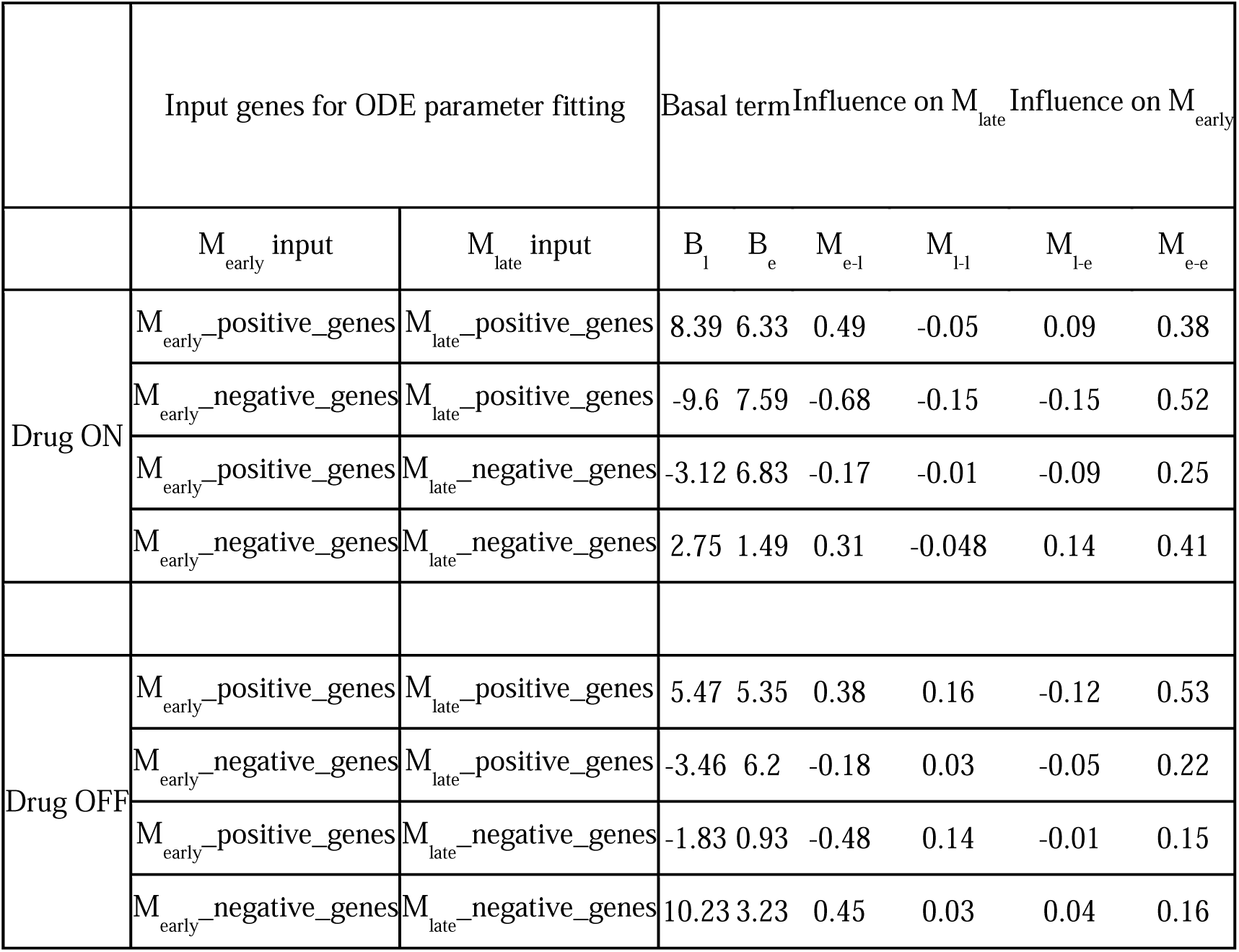
Best fitted parameters used in the dynamic ODE modeling.

